# Optogenetic stimulation of septal somatostatin neurons disrupts locomotory behavior and regulates hippocampus cholinergic theta oscillations

**DOI:** 10.1101/2022.09.30.510330

**Authors:** Nelson Espinosa, Mauricio Caneo, Alejandra Alonso, Constanza Moran, Pablo Fuentealba

## Abstract

The septal complex regulates both motivated and innate behaviors, chiefly by the action of its diverse population of long-range projection neurons. Among those cells are lateral septum somatostatin neurons which collateral axons profusely innervate cortically-projecting neurons located in the medial septum. Thus, somatostatin cells are ideally positioned at the crossroads of ascending and descending modulatory pathways, likely supporting functional roles in both anatomical directions. Here, we used optogenetic stimulation and extracellular recordings in acutely anesthetized transgenic mice to show that septal somatostatin neurons can disinhibit the cholinergic septo-hippocampal pathway, thus enhancing the amplitude and synchrony of theta oscillations, while depressing sharp wave ripple episodes in the dorsal hippocampus. Nonetheless, photosuppressing septal somatostatin cells hindered goal-directed behavior in a spatial memory task by disrupting task engagement, evidenced in increased immobility, followed by repetitive self-grooming, a hallmark innate behavior. These results suggest that septal somatostatin cells can recruit ascending cholinergic pathways to promote hippocampal theta oscillations, while gating repetitive displacement behaviors mediated by descending subcortical pathways.

**Significance Statement:** A small population of somatostatin-expressing GABAergic cells in the lateral septum projects deep into subcortical regions, yet on its way it also targets neighboring medial septum neurons that profusely innervate cortical targets. We show here that selective inhibition of septal somatostatin cells exerts significant consequences in both ascending and descending synaptic pathways. Indeed, cortical targets increased the expression of hippocampal theta oscillations, which are relevant for sensorimotor processing, temporal coding, and a marker of anxious behaviour; whereas subcortical targets triggered repetitive self-grooming, an innate displacement behaviour. Our results suggest that septal somatostatin cells are a potential target for the control of altered innate behaviors in translational neuroscience.

## Introduction

Large-amplitude theta oscillations dominate field potential hippocampal activity during active locomotion (Vanderwolf, 1969) and cognitive processing (Buzsáki and Moser, 2013), but also during aroused immobility (Robinson, 1980) and anxiety (Gray and McNaughton, 2008). The theta rhythm has been proposed to be essential for sensorimotor integration, temporal coding, and synaptic plasticity (Gyorgy, 2002). Nonetheless, theta activity is not homogeneous, and two bands have been identified. Indeed, type 1 theta activity is fast (6-12 Hz), atropine-resistant, arises during voluntary or goal-directed motor behavior, whereas type 2 theta activity occurs during immobility, is slower (4-8 Hz), anesthetic-resistant, and sensitive to cholinergic-antagonists (Kramis et al., 1975; Sainsbury et al., 1987). While type 1 theta activity seems to depend on inputs from the entorhinal cortex and is associated with spatial encoding and novelty detection, type 2 theta activity requires functional afferents from the CA3 area and is associated with arousal and anxiety (Buzsáki, 2002). Importantly, both theta bands rely on the integrity of the septal complex, in particular the medial septum, which densely innervates the hippocampus and entrains the theta rhythm (Tsanov, 2017).

The septal complex is heavily interconnected with the hippocampus (Unal et al., 2015). While the medial septum provides ascending projections to the hippocampus and prominently contributes to the coordination of theta oscillations and locomotor behavior (Nita et al., 2007; Kang et al., 2017a), the lateral septum receives descending hippocampal projections and is also essential to regulate locomotion speed and innate behaviors (Bender et al., 2015). The main septo-hippocampal projection pathways are GABAergic (Freund and Antal, 1988), cholinergic (Frotscher and Léránth, 1985), and glutamatergic (Köhler et al., 1984). Ever since long-range projection GABAergic cells were described, it has been proposed that individual cells exert simultaneous local and distal inhibition (Jinno et al., 2007), thus suggesting that this type of neuron is ideally suited to synchronize local and remote circuits. In fact, in the basal forebrain somatostatin cells provide functional inhibitory input to all three long-range neuronal populations (Xu et al., 2015) and previous work has shown their relevance to regulate excitability in executive centers, like the prefrontal cortex (Espinosa et al., 2019b).

Consequently, here we tested the hypothesis that the functional integrity of septal somatostatin cells is required to sustain the hippocampal theta rhythm. Accordingly, we used optogenetic inhibition of somatostatin cells in the dorsal septum and found that it locally enhanced the septal spiking output, resulting in increased amplitude and synchrony of theta oscillations in the hippocampus. The effect was evident under anesthesia and sensitive to cholinergic antagonists locally applied to the hippocampus, suggesting that, at a minimum, type 2 theta activity was affected by somatostatin cells. Moreover, optogenetic suppression of somatostatin cells in chronically implanted animals strongly impaired goal-directed behavior in a spatial navigation task. Instead, it was followed by reduced mobility and repetitive self-grooming, a prominent innate displacement behavior. We conclude that somatostatin cells can regulate both ascending and descending septal pathways to control cortical oscillations and innate behaviors.

## Methods

All procedures involving experimental animals were performed in accordance with ARRIVE guidelines, reviewed and approved by university and national bioethics committees. Experiments were carried out with 8- to 30-week-old mice (n = 24 from either sex), in accordance with the institutional Ethics Committee (protocol ID 151027003).

### Animals

Three mice strains from Jackson laboratories (www.jax.org) were used in this study, C57Bl/6J (stock N° 000664), Ai39 (stock N° 014539, B6, 129S-Gt(ROSA)26Sortm39(CAG-HOP/EYFP)Hze/J), and Sst-IRES-Cre (stock N° 013044, Ssttm2.1(cre)Zjh/J and stock N° 018973, B6N.Cg-Ssttm2.1(cre)Zjh/J). They were used as controls and we refer to them as (Natronomonas pharaonis halorhodopsin) NpHR-animals in the text. Double transgenic animals were obtained from the breeding of Sst-IRES-Cre and Ai39 mice, so that they expressed functional (NpHR) exclusively in somatostatin cells. We refer to such animals as NpHR+ in the text. Mice were genotyped by PCR on ear biopsies using the primers: GGG CCA GGA GTT AAG GAA GA (common), TCT GAA AGA CTT GCG TTT GG (wild type forward), TGG TTT GTC CAA ACT CAT CAA (mutant forward) for CRE Mice, and CTT TAA GCC TGC CCA GAA GA (wild type reverse), ATA TCC TGC TGG TGG AGT GG (mutant forward), GCC ACG ATA TCC AGG AAA GA (mutant reverse), TCC CAA AGT CGC TCT GAG (wild type forward) from Integrated DNA Technologies.

### In vivo electrophysiological recordings

Animals were induced with isoflurane, and then anesthetized with urethane (0.8 g/kg), and after 20 minutes a dose of ketamine (40 g/kg)/xylazine (4 g/kg) to start the surgical procedures. Throughout the experiment 1/12 of the initial dose of urethane was administered every 20–30 minutes. All drugs were administered intraperitoneally. Rectal temperature was monitored throughout the experiment and was kept at 36 ± 1 °C with a heating pad. Glucosaline solution was injected subcutaneously every 2 hours. In fully anesthetized mice, the scalp was cut and retracted to expose the skull. Mice were then implanted with a customized lightweight metal head holder and the head was held in a custom-made metallic holder. Next, small craniotomies (∼1 mm) were made with a dental drill above the basal forebrain, to target medial septum (MS; AP 1.0 mm, ML 0.0 mm, DV 2.5 mm from Bregma) and the hippocampus, to target the CA1 region (AP -2.5 mm at an angle of 20°, ML 2 mm, DV 1.2 mm from Bregma). Neuronal activity in MS was recorded by using a 16-channel silicon probe (A1 × 16-Poly2-Std, Neuronexus) stained with DiI and connected to an optic fiber (200 μm in diameter) attached to the shank (optrode), so electrical recording and photostimulation could be achieved simultaneously on the same site. Neuronal activity in the hippocampus was recorded extracellularly with a 32 channel-4 shank silicon probe (Buzsáki 32, Neuronexus) (mean resistance 1MΩ) stained with DiI. Electrical activity was recorded with a 32-channel Intan RHD 2132 amplifier board connected to an RHD2000 evaluation system (Intan Technologies). Single-unit activity and local field potential (LFP; sampling rate 20 kHz) were digitally filtered between 300 Hz–5 kHz and 0.3 Hz–2 kHz, respectively. Spike shape and amplitude were monitored during recording to ensure that the same cells were recorded. To allow local drug injection the hippocampal craniotomy was extended in the ML direction and a 50 μm-tip pipette was inserted dorso-ventrally with a 20° angle towards the midline. For the blockade of cholinergic receptors in CA1 200nl of atropine (2 mM) and mecamylamine (2 mM) (1:1) (Sigma Aldrich) were microinjected at 1.2mm DV (IM-9B microinjector, Narishige), at minute 5 of the recording, while giving pulses of light on the MS and recording from hippocampus.

### Surgery for Chronic Implantation

Mice were anaesthetized with isoflurane (4% induction and 1.5–2% maintenance) and placed on a stereotaxic frame (David Kopf Instruments). Temperature was kept at 37° throughout the procedure (1–2 h) using a heating pad. The skin was incised to expose the skull and a craniotomy (∼1mm in diameter) was made with a dental drill above the MS of the basal forebrain (AP + 0.55 mm and ML 0 mm from Bregma) (Franklin and Paxinos 2007). An optic fiber (diameter 200 um, length 11 mm; Thorlabs) inserted and glued to ceramic ferrules (diameter 230 um, length 6.4 mm; Thorlabs) were descended through the craniotomy until reaching the MS (DV 2 – 2.2 mm) and anchored to the skull using dental cement. After surgery, mice received a daily dose of enrofloxacin (10 mg/kg, Centrovet) for 5 days and supplementary analgesia with ketoprofen (5 mg/kg, Centrovet) for 3 days. Animals were allowed at least a week of recovery before behavioral tests.

### Optogenetic Stimulation

Optogenetic stimulation of somatostatin neurons was achieved with a 200 μm optic fiber, (N.A. 0.37, Thorlabs) coupled to a green laser (532 nm) that provided a total light power of 0.1–60 mW at the optrode tip. Light stimuli consisted of 5 second light pulses and power at the tip of the fiber was set at 10–15 mW for 200 μm optic fiber.

For anesthetized animals, every recording session lasted 10 minutes, during which laser stimulation was continuously presented for 5 seconds every 20 seconds (i.e., 15 sec. off – 5 sec. on). For chronically implanted animals, a 5 second light pulse was presented at the beginning of each trial.

### T-maze

Working memory was tested by recording the animals’ performance in an appetitive T-maze task. The apparatus consisted of two perpendicular blocks of black painted acrylic (central and lateral arms, length = 70 cm, width = 15 cm, height = 20 cm). The maze was equipped with three manually removable doors (15 × 20 cm). One door separated a 10-cm compartment at the beginning of the central arm (start). The other two doors were placed at the entrance of each lateral arm and blocked access to the lateral arms. The apparatus was placed in a testing room and illuminated by a 60 W bulb located 150 cm above.

For training, mice were food deprived until reaching 85% of their normal body weight. Animals were habituated on the T-maze with food (6-8 pieces of reward spread throughout the maze), for 2 days and 2 min sessions each day. After habituation, reinforced alternation training was initiated. Mice were trained in pairs of trials. On the first (forced) trial of the pair, a door blocked one of the lateral arms of the maze, while the unblocked lateral arm was baited with a small piece (10 mg) of sweetened cereal, placed 5 cm away from the arm’s end (reward). The animal was placed at the start position in the central arm at the beginning of every trial, after consuming the reward, animals were returned to the start position and held in place by a door blocking the central arm. For the second trial (choice) of the pair, the blocked lateral arm was opened and baited with reward. After 15 s, the blocking door was lifted. A choice was considered ‘correct’ when the animal entered the baited lateral arm. Conversely, ‘incorrect’ choices were punished with a 30 s timeout. Finally, a ‘null’ choice was considered when the animal either stayed in the central arm or entered to the baited arm without reaching or consuming the reward within 90 s. Choice trials were not performed when forced trials were null. The baited lateral arm in any given trial pair was pseudo-randomly assigned. Mice were tested in 8 pairs of trials per day (one session, 16 trials for sessions without null trials) for 8 consecutive days and with 2 min intervals between pairs during which time mice were placed in a plastic retention box similar to their home cage (19 × 30 × 12 cm). The T-maze was thoroughly cleaned with ethanol spray (10%) between trials and animals. The test was recorded using a digital video camera with a frame rate of 30 FPS.

### Histology

At the end of recordings, mice were terminally anesthetized and intracardially perfused with saline followed by 20 min fixation with 4% paraformaldehyde. Brains were extracted and postfixed in paraformaldehyde for a minimum of 8 h before being transferred to PBS azide and sectioned coronally (60–70 μm slice thickness). Sections were further stained for Nissl substance. Location of shanks and optical fiber were determined referred to standard brain atlas coordinates under a light transmission microscope.

### Behavioral Event Analysis

Test videos were analyzed manually on a frame-by-frame basis using a custom Matlab (MathWorks, Natick, MA) script to score performance (correct, incorrect, and null trials), latencies and grooming intervals. The investigator observer performing annotation was blind to experimental conditions. Additionally, animal position was tracked in Python with the open source toolbox DeepLabCut (Mathis et al., 2018), which provides markerless tracking of visual features in a video.

### Spike sorting

Semiautomatic clustering was performed by KlustaKwik, a custom program written in C++ (https://github.com/kwikteam/klustakwik2/). This method was applied over the 32 channels of the silicon probe, grouped in eight pseudo-tetrodes of four nearby channels. Spike clusters were considered single units if their auto-correlograms had a 2-ms refractory period and their cross-correlograms with other clusters did not have sharp peaks within 2 ms of 0 lag.

### Unit cross-correlation analysis

Neural activity in the MS and hippocampus was cross-correlated with the light pulse by applying the “sliding-sweeps” algorithm. A time window of ±15 s was defined with point 0 assigned to the light onset. The timestamps of the hippocampal and basal forebrain spikes within the time window were considered as a template and were represented by a vector of spikes relative to t = 0 s, with a time bin of 500 ms and normalized to the basal firing rate of the neurons. Thus, the central bin of the vector contained the ratio between the number of neural spikes elicited between ± 250 ms and the total number of spikes within the template. Next, the window was shifted to successive light pulses throughout the recording session, and an array of recurrences of templates was obtained. Both neural timestamps and start times of light pulses were shuffled by randomized exchange of the original inter-event intervals and the cross-correlation procedure was performed on the random sequence.

### Spectral analysis

Time-frequency decomposition of LFP was performed with multi-taper Fourier analysis implemented in Chronux toolbox (http://www.chronux.org). LFP was downsampled to 500 Hz before decomposition. The same taper parameters described for the coherence analysis were used. To estimate gamma band power, spectra were normalized by 1/f to correct for the power law governing the distribution of EEG signals. To compute power and frequency of the gamma band oscillation, LFP was band-pass filtered with a two-way least squares finite impulse response (FIR) filter (eegfilt.m from EEGLAB toolbox; http://www.sccn.ucsd.edu/eeglab/).

### Ripples detection

Sharp wave-ripples were recorded in dorsal CA1, as close as possible to stratum pyramidale. We used a previously described method for ripples detection (Logothetis et al. 2012) with some variation. Briefly, the hippocampus LFP was first down-sampled to 500 Hz, then band-pass filtered (100-250 Hz) using a zero-phase shift non-causal FIR filter with 0.5 Hz roll-off. Next, the signal was rectified, and low-pass filtered at 20 Hz with a 4th order Butterworth filter. This procedure yields a smooth envelope of the filtered signal, which was then z-score normalized using the mean and SD of the whole signal in the time domain. Epochs during which the normalized signal exceed a 3.5 SD threshold were considered as ripple events. The first point before threshold that reached 1 SD was considered the onset and the first one after threshold to achieve 1 SD as the end of events. The difference between onset and end of events was used to estimate the ripple duration. We introduced a 50 ms-refractory window to prevent double detections. To precisely determine the mean frequency, amplitude, and duration of each event, we performed a spectral analysis using Morlet complex wavelets of seven cycles. The Matlab toolbox used is available online as LAN toolbox (https://bitbucket.org/marcelostockle/lan-toolbox/wiki/Home).

### Coherency

Both spike-field and LFP-LFP coherence were computed using the multitaper Fourier analysis and the Chronux toolbox (http://www.chronux.org). We used 500 data points at 1000 Hz, a time-bandwidth product (TW) of 3 and 5 tapers, resulting in a half width of 0.6 Hz.

### Pairwise phase consistency

Pairwise phase consistency (PPC, (Vinck et al., 2010)) was computed with ft_connectivity_ppc.m, a Matlab function implemented in Fieldtrip (http://www.fieldtriptoolbox.org/reference/ft_connectivity_ppc). Briefly, the phase was extracted using Hilbert transform and the mean of the cosine of the absolute angular distance among all pairs of phases was calculated. This procedure was applied from 0 to 10 Hz (bin = 0.25 Hz).

### Granger causality

The Multivariate Granger causality (MVGC) Matlab toolbox (Barnett and Seth, 2014) was used to assess pairwise causalities between LFP-LFP and LFP-multiunit activity. This toolbox, available online (https://users.sussex.ac.uk/~lionelb/MVGC/html/mvgchelp.html#1), allows a fast and accurate estimation of the Wiener-Granger causal inference in the frequency domain. Estimator were calculated with the standard ordinary least squares and with a model order of 50. Frequency resolution was set at 1000.

### Statistics

Data sets were tested for normality using Kolmogorov-Smirnov test and then compared with the appropriate test (t-test or Wilcoxon two-sided rank sum test). Statistical significance of data for protocols with factorial design (i.e., light on/off conditions) were assessed using two-way repeated-measures ANOVA followed by false discovery rate (FDR) for multiple comparison corrections or Kruskal Wallis test followed by a Mann-Whitney U contrasts.

## Results

### Septal somatostatin cells regulate firing patterns in the medial septum and dorsal hippocampus

We used transgenic mice selectively expressing YFP-labeled halorhodopsin (NpHR) in somatostatin cells. We first established the relevance of somatostatin cells for the spontaneous activity patterns in the medial septum. For this, we stereotaxically implanted a silicon probe in the medial septum of urethane-anesthetized transgenic animals, coupled with an optic fiber on the dorsal septum (**Fig. 1A, Fig.1-1**). As previously reported, we used extended laser pulses to achieve maximal inhibition of somatostatin cells and reproduce prior experimental protocols (Xu et al., 2015; Espinosa et al., 2019a, 2019b). In doing so, we obtained diverse types of response in individual septal units (**Fig. 1B**), yet the net effect of optogenetic stimulation on the medial septum was excitation, as the global firing rate increased by 12.7% on average (**Fig. 1C**). Indeed, upon optogenetic stimulation a small fraction of septal cells (7.3%, n = 13 units) strongly decreased its activity (by 45.5 ± 4.3%), whereas another minor neuronal population (21.5%, n = 38 units) increased its firing rate (by 53.2 ± 7.9 %), presumably by synaptic disinhibition (**Fig. 1D, Fig. 1-2**). Baseline firing rates and spike waveforms were very similar between excited and inhibited units (**Fig. 1-2**), consistent with previous results in the basal forebrain (Espinosa et al., 2019b). Next, we estimated whether the increased neuronal discharge was associated to specific frequency bands of activity. For this, we computed the power of neuronal activity across the frequency spectrum. A fraction of medial septum units exhibited preference to discharge in the low theta frequency band (2-6 Hz). Interestingly, such tendency was selectively abolished upon optogenetic suppression of somatostatin cells (P = 0.016, **Fig. 1E**). We also assessed the effect of the medial septum’s synaptic output onto dorsal hippocampal targets. Indeed, our medial septum recordings were coupled with simultaneous monitoring of dorsal hippocampal activity (**Fig. 1A**), where single units exhibited mixed results upon optogenetic stimulation in the dorsal septum (**Fig. 1B**). The overall effect of dorsal septum optogenetic stimulation in the dorsal hippocampus was excitation, as the global firing rate increased by 8.4% (**Fig. 1F**). Inhibition of septal somatostatin cells robustly affected hippocampal spiking activity, with a significant proportion of hippocampal neurons (20.2%, n = 65) consistently increasing their firing rate (by 34.1 ± 3.4%), whereas a minor proportion of hippocampal units (4.7%, n = 15) was inhibited (by 19.4 ± 3.3%) by laser stimulation. Laser-excited hippocampal units had low spontaneous firing rates, broad spike waveforms and were strongly coupled to sharp-wave ripples (**Fig. 1-2**), consistent with the characteristics of pyramidal cells (Csicsvari et al., 1999). Conversely, laser-inhibited hippocampal cells showed high spontaneous firing rates, narrow spikes, and were moderately coupled to sharp-wave ripples (**Fig. 1-2**), consistent with the characteristics of interneurons (Csicsvari et al., 1999). Thus, our results suggest that optogenetic suppression of septal somatostatin neurons disinhibits pyramidal cells in the hippocampus, probably by decreasing interneuron activity (**Fig. 1G**). Spiking activity of hippocampal units exhibited bimodal preference in the frequency spectrum, with prominent peaks in the theta band (3-7 Hz) and the fast-frequency range (200 Hz, **Fig. 1H**), likely resulting from bursting discharges of pyramidal cells (**Fig. 1-2**). Interestingly, optogenetic inhibition of septum somatostatin cells selectively increased the amplitude of theta-frequency discharge in hippocampal units, without affecting the fast-frequency range (P = 3×10^−4^, **Fig. 1H**). Thus, septum somatostatin cells can regulate the activity of local and distal synaptic targets, differentially affecting neuronal firing rates and spectral frequency preferences.

**Figure 1.**
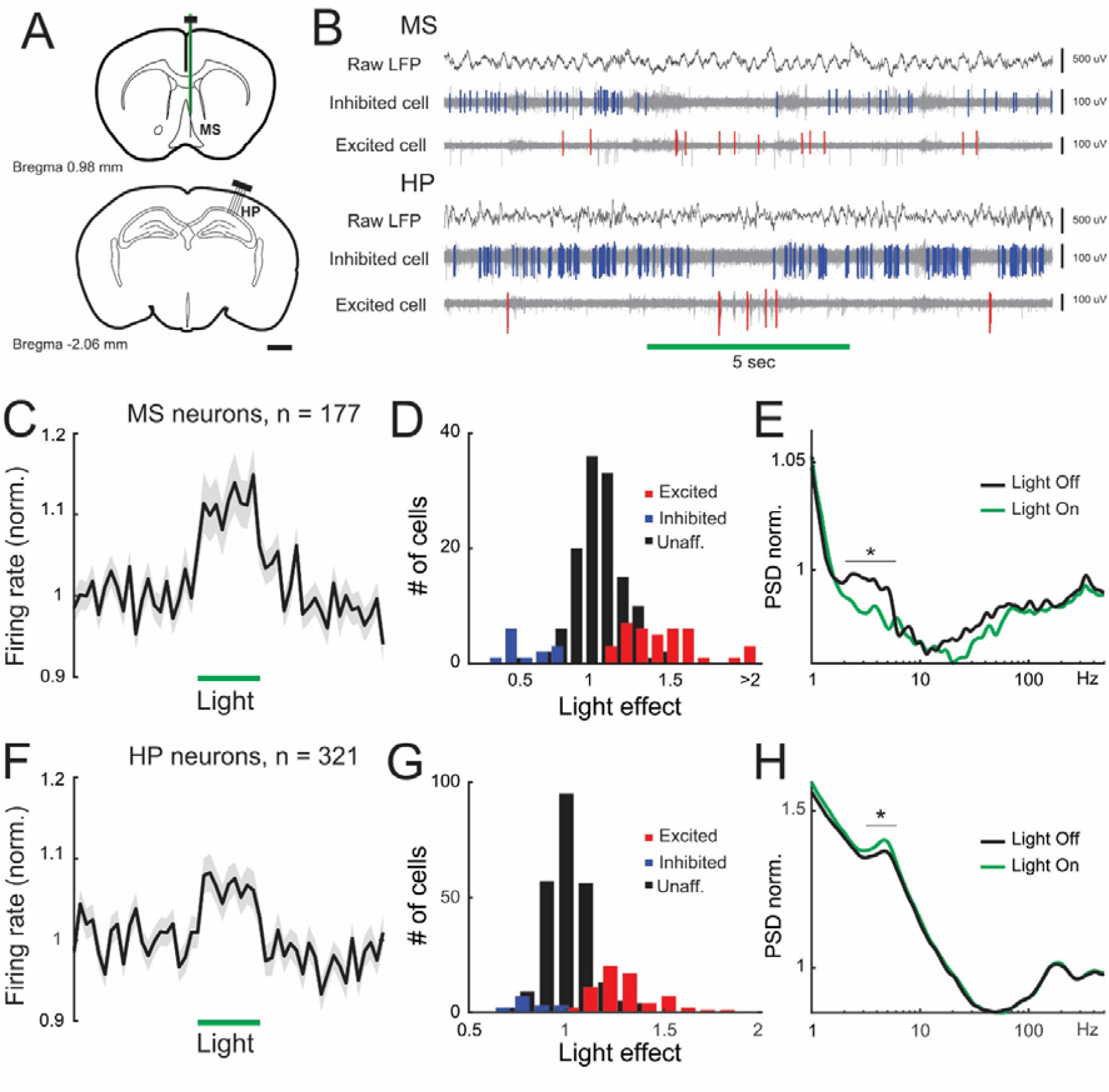
Neuronal activity in the medial septum and dorsal hippocampus during optical inactivation of somatostatin cells in the medial septum. **A**, recording locations represented in schematic coronal brain sections. A four-shank multielectrode angled 20° from the midline and an optrode (vertical line) were stereotaxically implanted in the dorsal hippocampus and medial septum, respectively. **B**, example simultaneous electrophysiological recordings (mouse NE115, recording 01) showing LFP and multiunit activity (LFP filtered 500 Hz–5 kHz) from the medial septum (MS, channels 39, 48, and 34 respectively) and dorsal hippocampus (HP, channels 20, 16, and 25 respectively). Horizontal bar depicts laser stimulation (5 s, 10 mW fiber diameter 200 um) and colors depict inhibited (blue events) and excited (red events) units. **C**, average normalized discharge probability for all septal neurons (mean value ± SEM, n = 177 units, 7 animals). Bin size: 500 ms. **D**, mean firing rate (z-score) distribution during light stimulation for septal neurons. Light and dark grey bars denote statistical difference from shuffled data. **E**, averaged power spectral density (PSD) for all recorded neurons before (black line) and during light stimulation (grey line). Asterisk in band 2 – 6 Hz: paired t-test, P = 0.016. **F**, average normalized discharge probability for all hippocampal neurons (mean value ± SEM, n = 321 units, 7 animals). Bin size: 500 ms. **G**, mean firing rate (z-score) distribution during septal stimulation for hippocampal neurons. Light and dark grey bars denote statistical difference from shuffled data. **H**, averaged power spectral density (PSD) for all recorded neurons before (black line) and during light stimulation (grey line). Asterisk in 4 – 7 Hz: paired t-test, P = 0.0003. Supported by Extended Data Figures 1-1 and 1-2.

### Inhibition of septal somatostatin cells enhances theta oscillations in the dorsal hippocampus

We then studied the field potential effects of septal optogenetic stimulation (**Fig. 2**). Inhibition of somatostatin cells did not show evident changes in the local field potential of the medial septum (**Fig. 2-1**). Conversely, optogenetic stimulation selectively increased the power of theta oscillations in the dorsal hippocampus (**Fig. 2A**) by 14.8% (P = 8×10^−7^, **Fig. 2B**), with no noticeable effects on other prominent rhythms such as delta or gamma oscillations (**Fig. 2B, Table 2-1**). Intrahippocampal synchrony of theta oscillations was also augmented during optogenetic stimulation, as evidenced by the increased coherence of field potentials, particularly for theta oscillations (P = 1.6×10^−4^, **Fig. 2C**). Further, hippocampal cells increased their coupling with theta oscillations during optogenetic stimulation, as evidenced by both enhanced spike-field coherence (P = 9.9×10^−8^, **Fig. 2D**) and pairwise phase consistency (**Fig. 2-2**). In addition, silencing somatostatin cells significantly decreased the density of sharp-wave ripples (**Fig. 2E**) by 17.3% (P = 0.013, **Fig. 2F**). These results are consistent with the selective activation of ascending cholinergic transmission from the medial septum (Vandecasteele et al., 2014; Dannenberg et al., 2015). To further test this idea, we locally applied cholinergic receptor antagonists in the dorsal hippocampus during optogenetic inhibition of septal somatostatin cells (**Fig. 2G**). Cholinergic blockers transiently abolished the enhancement of hippocampal theta oscillations (**Fig. 2H**), an effect specific for that frequency band (P = 0.259), as it did not affect other cortical rhythms (**Fig. 2H, Table 2-2**). While these experiments do not exclude the involvement of GABAergic and glutamatergic transmission, they suggest that enhanced hippocampal theta oscillations during optogenetic inhibition of septal somatostatin cells were largely mediated by cholinergic transmission in the hippocampus

**Figure 2.**
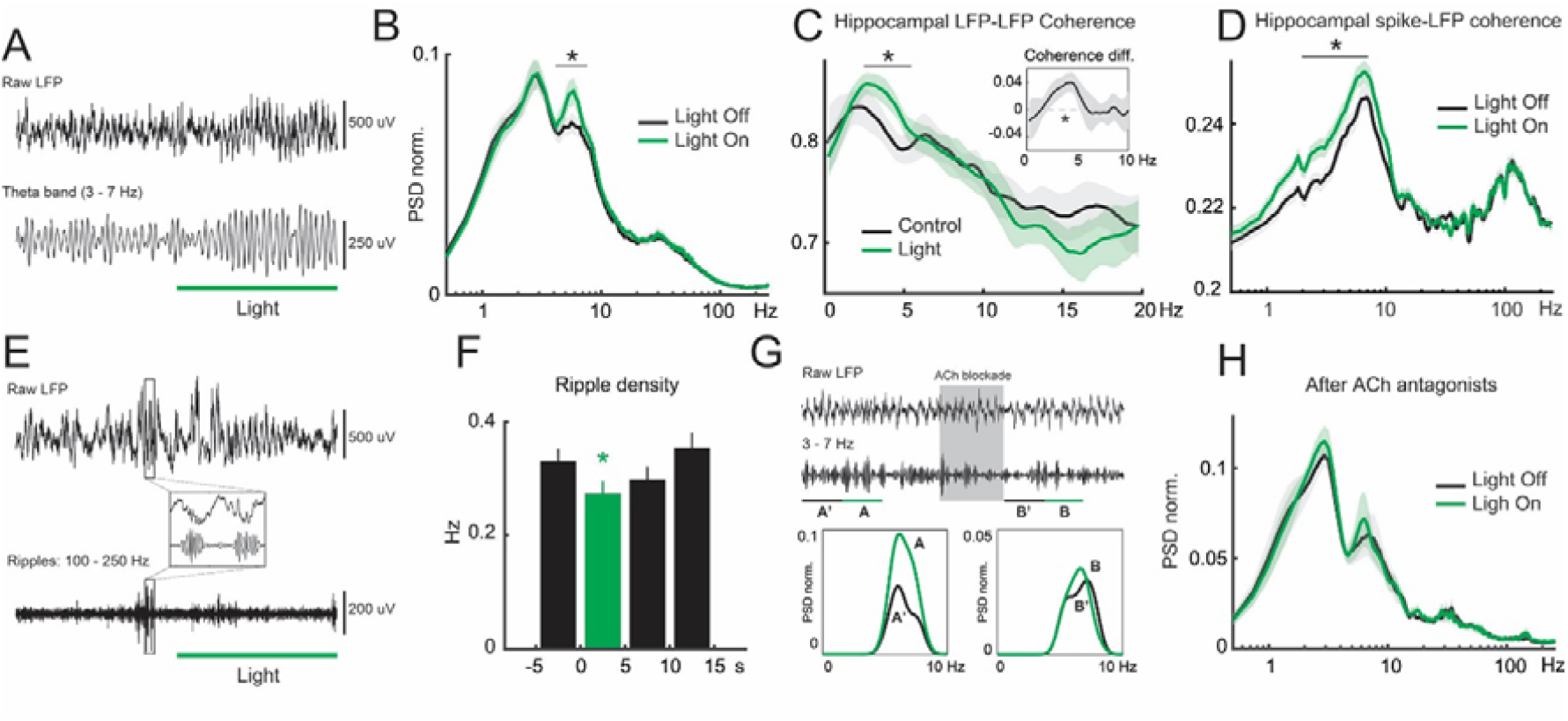
Theta oscillations and ripple density in the dorsal hippocampus during septal somatostatin cells inhibition. **A**, raw LFP (top panel), and theta-band filtered LFP (3 – 7 Hz, bottom panel) for an example electrophysiological recording in the dorsal hippocampus (mouse NE113, recording 05, channel 29). **B**, average normalized power spectral density (PSD) of the dorsal hippocampal LFP in the presence (grey line) or absence (black line) of optogenetic stimulation. Asterisk in band 4 – 7 Hz: paired t-test, P = 7.99×10^−7^, n = 27 recordings. **C**, average LFP-LFP coherence around the pyramidal layer (i.e., between the more dorsal and ventral channels of the shank). Asterisk in band 3 – 6 Hz: paired t-test, P = 0.00016, n = 27 recordings. Inset, mean difference between spectral distributions (Control - Light). **D**, average spike-field coherence between dorsal hippocampal units and LFP pyramidal layer. Asterisk in band 2 – 7 Hz: W = 16,976, P = 9.9×10^−8^, Wilcoxon signed-rank test, n = 321 cells. **E**, raw LFP (top panel), and ripple-band LFP (100 – 250 Hz, bottom panel) for an example electrophysiological recording in the dorsal hippocampus (mouse NE113, recording 05, channel 29). Inset, sharp wave ripple episodes before light stimulation. **F**, septal somatostatin cells inhibition decreases ripple density (grey bar, mean value ± SEM, bin = 5 s). Asterisk: one-way ANOVA test, and Bonferroni post-hoc correction, P = 0.0131, n = 262 trials. **G**, example electrophysiological recording (mouse NE125, recording 03, channel 20) showing response patterns before and after drug administration (gray box). From top to bottom: raw LFP, theta-band filtered LFP, and PSD during control (black line) and optogenetic stimulation (grey line). **H**, average normalized power spectral density (PSD) of the dorsal hippocampal LFP after cholinergic receptors blockade. Supported by Extended Data Figures 2-1 and 2-2.

Next, we investigated the directionality of the functional relationship between the medial septum and dorsal hippocampus. To assess directionality, we used Granger causality within the theta frequency band (Kang et al., 2017a). We found significant, unidirectional connectivity from the septum to the hippocampus (MS->HP) in the low theta frequency band (3-6 Hz, **Fig. 3A**). Importantly, the MS->HP connectivity was strongly dependent on the activity of somatostatin cells, as their optogenetic suppression significantly reduced the causality (P = 2.4×10^−6^, **Fig. 3A**). Reduction in Granger causality was specific to the theta band, as it did not affect other rhythms, such as gamma oscillations (**Fig. 3A, Table 3-1**). Moreover, the reciprocal connectivity, HP->MS, did not show apparent relevance for theta band synchrony, neither was it dependent on the activity of septum somatostatin neurons (P = 0.078, **Fig. 3B**). We further analyzed multiunit activity from the medial septum and found similar patterns in which unidirectional connectivity from the medial septum to the dorsal hippocampus was dependent on the activity of somatostatin cells (**Fig. 3-1**). Previous reports have shown that as theta power increases during activated brain states, and complementary delta waves decrease, theta Granger causality in the HP->MS direction selectively increases, but not in the reverse MS->HP direction (Kang et al., 2017b). Hence, we tested whether the amplitude of slow oscillations was inversely proportional to theta causality in the HP->MS direction in our recordings. We found no significant correlation during baseline conditions (P = 0.08; **Fig. 3C**), yet upon optogenetic suppression of somatostatin cells it became apparent (Spearman’s R = -0.17, P = 8×10^−4^; **Fig. 3D**). Indeed, high power in hippocampal delta waves was associated with low Granger causality in the theta band (**Fig. 3E**), whereas low power in the delta frequency band was related to statistically significant theta Granger causality (**Fig. 3F**). Hence, suppressing somatostatin cells enhance cholinergic theta oscillations in the dorsal hippocampus and regulate the directionality of connectivity between medial septum and dorsal hippocampus.

**Figure 3.**
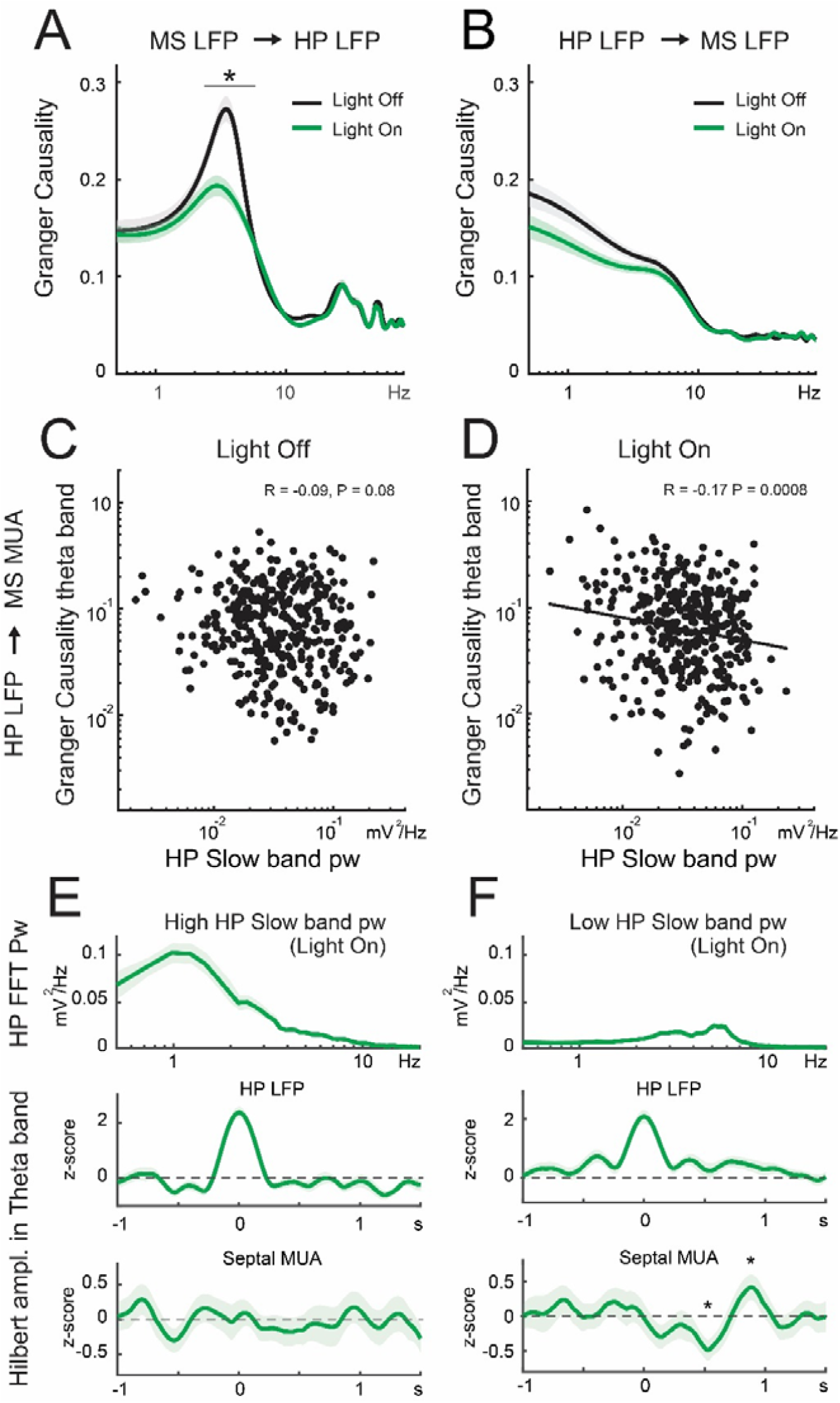
Granger causality between medial septum and dorsal hippocampus during optical inactivation of septal somatostatin cells. **A**, averaged Granger causality from the septal LFP to hippocampal LFP (MS LFP -> HP LFP). Septal somatostatin cells inhibition statistically decreases the peak frequency. Asterisk in band 3 – 6 Hz: W = 52,226, P = 2.4×10^−6^, Wilcoxon signed-rank test, n = 405 trials. **B**, reverse Granger causality (HP LF -> MS LFP) is unaffected by light stimulation. **C**, under control condition (light off) Granger causality from the hippocampal LFP to septal multiunit activity (HP LFP -> MS MUA) is uncorrelated with the hippocampal slow wave power. **D**, significant correlation between Granger causality HP LFP -> MS MUA emerges during light stimulation (Spearman’s r = -0.17, P = 0.0008, n = 381). Black line depicts the least-squares regression line fitting. **E**, average values for the 5^th^ lower percentile in Granger causality HP LFP -> MS MUA and 5^th^ upper percentile in hippocampal slow wave power (n = 19 trials) of trials during light stimulation (D). Top panel: average normalized power spectral density of hippocampal LFP. Middle panel: z-scored theta-filtered amplitude for hippocampal LFP averaged with peak value set at t = 0. Bottom panel: average z-scored theta-filtered amplitude for septal LFP and referred to peak value of hippocampal theta. **F**, average values for the 5^th^ upper percentile in Granger causality HP LFP -> MS MUA and 5^th^ lower percentile in hippocampal slow wave power (n = 19 trials) of trials during light stimulation (D). Top panel: average normalized power spectral density of hippocampal LFP. Middle panel: z-scored theta-filtered amplitude for hippocampal LFP averaged with peak value set at t = 0. Bottom panel: average z-scored theta-filtered amplitude for septal LFP and referred to peak value of hippocampal theta. Asterisk denotes significant difference against zero (Wilcoxon signed-rank test, FDR correction, P < 0.0142). Supported by Extended Data Figure 3-1.

### Inhibition of septal somatostatin cells disrupts goal-directed behavioral engagement

The medial septum is closely involved with the control of both theta oscillations and locomotion speed (Espinosa et al., 2019a; Lu et al., 2020). To assess the role of septum somatostatin cells on spatial navigation and locomotor behavior, we stereotaxically implanted optical fibers on the dorsal septum and trained mice to perform a goal-directed navigation task, the delayed non-matched to position task (**Fig. 4A**). We delivered laser stimulation at the start of both phases (sample and test) and assessed learning performance in both transgenic (NpHR+) and control (NpHR-) animals. After training, control mice significantly improved their performance (day 1 = 0.5 ± 0.09 vs. day 8 = 0.8 ± 0.02, P = 0.04, paired t-test; **Fig. 4-1**), whereas transgenic animals, expressing functional NpHR, did not exceed chance level (day 1 = 0.22 ± 0.07 vs. day 8 = 0.39 ± 0.06, P = 0.12, paired t-test; **Fig. 4-1**), revealing an impairment in task acquisition (**Fig. 4A**). Furthermore, control mice showed a progressive improvement in task acquisition over time (Spearman’s R = 0.34, P = 0.009), which was not detected in transgenic mice (P = 0.12). Not only performance, but also task latency was compromised during optogenetic stimulation as control mice progressively decreased the time to reach the target arm (day 1 = 25.2 ± 2.2 s vs. day 8 = 4.9 ± 0.3 s, P = 4.6 × 10^−12^, unpaired t-test), whereas transgenic animals exhibited consistently long delays (day 1 = 43.2 ± 3.0 s vs. day 8 = 42.5 ± 3.0 s, P = 0.88, unpaired t-test) during the entire experimental protocol (**Fig. 4B, Table 4-1**). The longer latency of transgenic animals was not the result of prolonged exploration of the T-maze. On the contrary, tracking individual trajectories of mice performing the spatial task allowed us to construct occupation maps, which revealed that while control animals did not show apparent preference to explore particular regions of the maze, transgenic animals remained immobile for long periods at the start area (4.5 ± 0.37 s for NpHR-vs. 35.4 ± 1.1 s for NpHR+; W = 756400, P < 10^−138^, Wilcoxon rank-sum test; **Fig. 4C**). Indeed, during the first days of training, both transgenic and control animals had the tendency to remain at the start of the maze, but from then onwards control mice started to explore the entire maze, whereas transgenic mice remained stationary at the start area (**Fig. 4-1**). Consistent with that observation, the traveled distance in the maze progressively diverged and became consistently different between control and transgenic mice as training advanced (**Fig. 4D, Table 4-1**). Indeed, control mice steadily covered more distance in the maze throughout training (Spearman’s R = 0.32, P < 10^−22^; least-squares regression slope = 4.3 cm/day, bootstrap corrected), while transgenic animals showed little variation in their spatial exploratory pattern (Spearman’s R = 0.04, P = 0.22, **Fig. 4D**).

**Figure 4.**
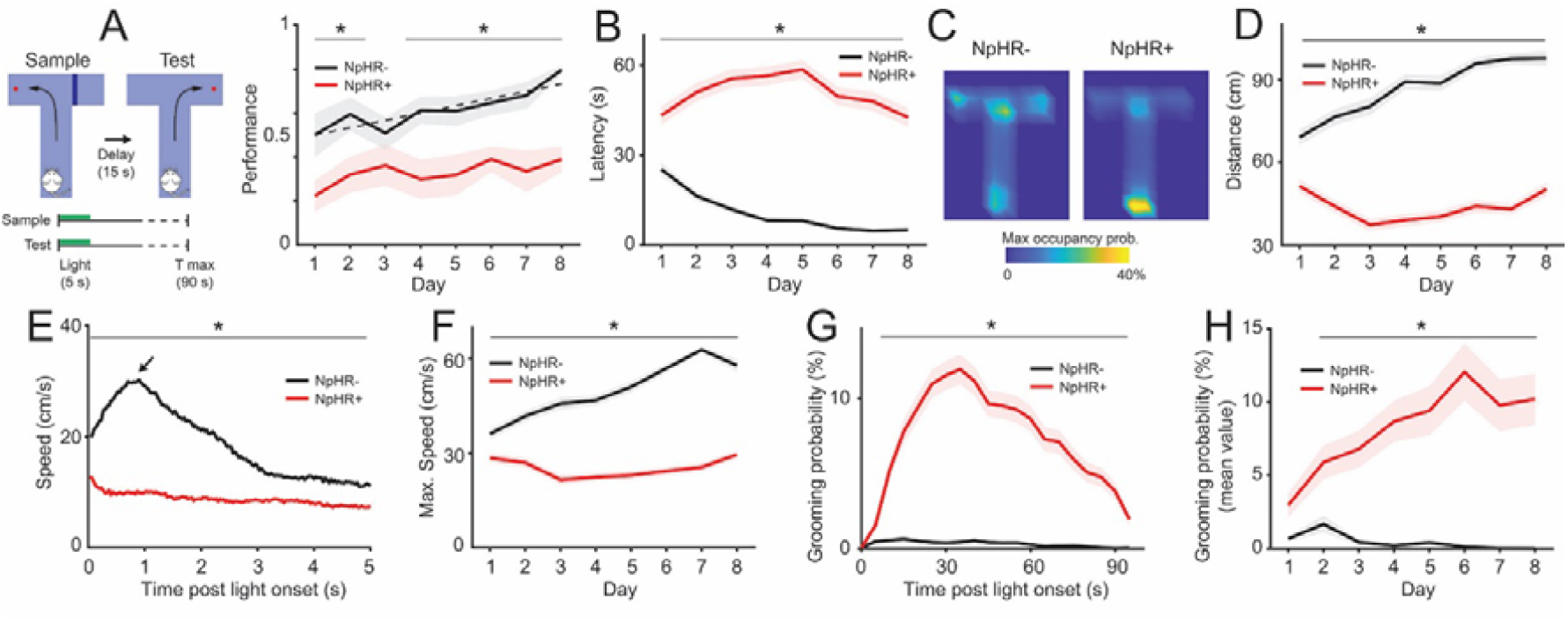
Inhibition of septal somatostatin cells during goal-directed behavior. **A**, left; schematic of the T-maze forced alternation task used for control (NpHR-) and transgenic (NpHR+) animals with a light pulse (5 s) delivered at the beginning of both sample and test phases. Right; performance across days for NpHR-(mean ± SEM, black line, Spearman’s r = 0.34, P = 0.009, 8 animals) and NpHR+ (mean ± SEM, grey line, Spearman’s r = 0.15, P = 0.12, 9 animals) groups. Dashed line depicts the corresponding least-squares linear regression for NpHR-animals. *P < 0.018, Wilcoxon rank sum test. **B**, task latency across days. *P < 10^−5^, Wilcoxon rank sum test. **C**, average normalized occupancy at each location within the T-maze. **D**, total traveled distance in the T-maze. *P < 10^−21^, Wilcoxon rank sum test. **E**, average instantaneous speed during light stimulation (first 5 s, bin = 33.3 ms). *P < 10^−8^, Wilcoxon rank sum test. **F**, average maximum instantaneous speed across days. *P < 3×10^−4^, Wilcoxon rank sum test. **G**, probability of grooming during trials. Bin = 5 s. Grey bar indicates light stimulation. *P < 4×10^−4^, Wilcoxon rank sum test. **H**, average probability of grooming across days. *P < 2×10^−6^, Wilcoxon rank sum test. (B) – (H): values in mean ± SEM; NpHR-, black line. NpHR+, grey line. N = (n-animals) x (n-choices), where n-choices depends on the number of null forced trials per day (i.e., when choice trial is omitted, see text for details). *P, significant pairwise comparison across days corrected by FDR. Supported by Extended Data Figures 4-1, 4-2, 4-3, and 4-4.

Next, we assessed the instantaneous speed of mice during navigation as an indicator of locomotion patterns. On average, control animals quickly increased their speed during the first second of exploration in the maze, and then rapidly decreased as they approached the maze junction (**Fig. 4E**). In fact, maximal locomotion speed gradually increased during training (Spearman’s R = 0.46, P < 10^−47^; least-squares regression slope = 3.6 cm/s*day, bootstrap corrected, **Fig. 4F, Table 4-1**), consistent with decreased latency and task acquisition. Conversely, transgenic animals showed low and stable speeds during the entire laser pulse (**Fig. 4E**), which did not evolve as training advanced (**Fig. 4-2**). Similarly, maximal speed remained steady along training days for transgenic mice (Spearman’s R = 0.03, P = 0.29, **Fig. 4F**). Overall, both mean and maximal locomotion speeds were consistently larger for control animals when compared to transgenic mice (**Fig. 4-3**). Importantly, transgenic mice were not intrinsically affected in their ambulatory or motor capacity as their locomotion speed (**Fig. 4-3**) and travelled distance (**Fig. 4-3**) were not different from control animals during early stages of the test (two-sample t-test, P = 0.11). Moreover, due to suppressed ambulatory behavior upon optogenetic stimulation, transgenic animals failed to perform a large proportion of experimental trials (33 ± 2.8%) when compared to control mice (1.7 ± 0.7%; P = 2.9×10^−21^, Wilcoxon rank sum test, W = 2635). To assess whether such suppression biased results from the population analysis, we omitted null trials to recalculate and compare behavioral parameters between animal groups. Hence, we confirmed that locomotor behavior was progressively compromised upon optogenetic stimulation, even when transgenic mice completed task trials (**Fig. 4-3**). Taken together, these results suggest that optogenetic suppression of septum somatostatin cells did not affect motor function, but impaired task acquisition by disrupting locomotor behavior.

Finally, we closely inspected mice behavioral patterns during episodes of reduced locomotion. Hence, we realized that transgenic mice exhibited repetitive self-grooming following laser stimulation. Indeed, transgenic mice rapidly increased the probability of self-grooming during the first 30 seconds in the maze, whereas control animals exhibited very low, persistent probability to perform grooming during the task trials (P = 5×10^−4^, **Fig. 4G**). Moreover, the mean probability of self-grooming in the maze increased during training days for transgenic animals (day 1 = 2.97 ± 0.79% vs. day 8 = 10.21 ± 1.73%, P = 2.8×10^−4^, Wilcoxon rank sum test), whereas it decreased for control mice throughout the behavioral protocol (day 1 = 0.68 ± 0.22% vs. day 8 = 0%, P = 6.7×10^−4^, Wilcoxon rank sum test, **Fig. 4H, Table 4-1**). Interestingly, self-grooming was much more likely during the sample phase of the task when compared to the testing phase, exclusively for transgenic animals (**Fig. 4-4**), suggesting short-term behavioral adaptation to laser stimulation. Furthermore, to establish baseline conditions of self-grooming in our experimental animals, we individually placed them in an open field (i.e., squared box) and allowed them to explore freely the environment while videorecording their behavior. Concurrently, we randomly delivered optogenetic stimuli to the dorsal septum and assessed self-grooming (**Fig. 4-4**). Under these conditions, we found that baseline probability of self-grooming was similarly stable for both experimental groups and not affected by optogenetic stimulation (**Fig. 4-4**). Thus, under baseline conditions self-grooming did not differ between experimental groups and, importantly, the apparent increase in self-grooming behavior of transgenic mice was highly sensitive to the behavioral context as it was specific for the navigation task. Overall, these results suggest that optogenetic suppression of septum somatostatin cells impaired goal-directed behavior by reducing normal locomotion and gating repetitive self-grooming.

## Discussion

We have shown that optogenetic inhibition of somatostatin cells in the dorsal septum locally enhances spiking output, including septo-hippocampal cells, thus resulting in increased amplitude of theta oscillations and decreased incidence of ripple episodes in the dorsal hippocampus. The effect involves the disinhibition of septal cholinergic pathways, as it was blocked by cholinergic antagonists locally applied to the hippocampus. Moreover, functional connectivity in the theta band was causal in the septo-hippocampal direction, and dependent on the activity of somatostatin cells. Finally, optogenetic suppression of somatostatin cells impaired goal-directed behavior navigation, and was followed by repetitive self-grooming, a prominent innate behavior. Our results show that somatostatin cells can regulate ascending septal pathways to promote cortical theta oscillations while also gating innate behaviors distinctively mediated by subcortical circuits.

In our experiments, laser inhibition of dorsal septum somatostatin cells produced net increased spiking activity in the medial septum, likely due to synaptic disinhibition (Ikeda and Wright, 1972). The neuroanatomical organization of the septal complex supports this interpretation as there are three main septo-hippocampal projection pathways conveyed by GABAergic (Freund and Antal, 1988), cholinergic (Frotscher and Léránth, 1985), and glutamatergic (Köhler et al., 1984) neurons; and basal forebrain somatostatin cells provide functional inhibitory input to all those neuronal populations (Zaborszky and Duque, 2000; Xu et al., 2015). Hence, those cell types are available to elevate their baseline activity levels, and become more sensitive to incoming synaptic barrages, upon optogenetic removal of the inhibitory drive supplied by somatostatin cells. Consequently, distal cortical targets, such as the dorsal hippocampus, were also activated after suppression of septal somatostatin cells.

In both medial septum and dorsal hippocampus, septal laser stimulation produced net excitation, as evidenced by the significant increase in overall firing rates. Nonetheless, the preferred spectral frequency of neuronal discharge was inversely influenced. Indeed, during laser stimulation hippocampal units increased their activity in the theta band, while septal units reduced their preference to discharge in the theta frequency. The theta discharge of medial septum cells was broader and slower than that of dorsal hippocampus units, consistent with the slow cholinergic theta activity associated to motor behavior originating from the subcortical nucleus, as compared to hippocampal theta oscillations, which also integrate faster noncholinergic components (Bland et al., 2006; Bland, 2008).

Direct optogenetic activation of septal cholinergic cells excites hippocampal interneurons, which in turn inhibit pyramidal cells (Dannenberg et al., 2015). Conversely, our septal laser stimulation seemed to produce opposite effects in hippocampal units, with enhanced putative pyramidal cell activity and decreased spiking in putative interneurons. This discrepancy may result from the differential connectivity between cholinergic cells and somatostatin neurons. For example, somatostatin cells are presynaptic to cholinergic neurons in the basal forebrain, and their respective activation produces opposite effects in the sleep-wake cycle (Xu et al., 2015). Similarly, silencing somatostatin cells or activating cholinergic cells in the septal complex seems to be sufficient to enhance theta oscillations in the dorsal hippocampus. Type 1 theta activity is faster (6-12 Hz) and arises during voluntary or goal-directed motor behavior, whereas type 2 theta activity is slower (4-8 Hz), occurs during immobility, is resistant to most anesthetics, and is sensitive to cholinergic-antagonists (Seidenbecher et al., 2003; Gordon et al., 2005; Gray and McNaughton, 2008; Adhikari et al., 2011). Hence, our evidence suggests that somatostatin cells participate, at least, in the regulation of hippocampal type 2 theta oscillations.

Further, spectral frequency preferences of individual unitary activity were mirrored in the field potential, as optogenetic inhibition of somatostatin cells significantly increased the power of theta oscillations in the dorsal hippocampus. The local application of cholinergic receptor antagonists confirmed that septo-hippocampal cholinergic pathways were largely responsible for the enhanced hippocampal theta waves (Lee et al., 1994). We have previously shown comparable results in the ascending basal forebrain projections innervating the medial prefrontal cortex (Espinosa et al., 2019b). Thus, we propose that the transient suppression of GABAergic input provided by somatostatin cells disinhibits septo-hippocampal cells, thus increasing the acetylcholine release on the dorsal hippocampus and consequently enhancing the power and synchrony of theta oscillations. Indeed, similar results were obtained in previous studies, using either patterned optogenetic stimulation of cholinergic septal neurons in vivo (Vandecasteele et al., 2014; Dannenberg et al., 2015) or bath applications of cholinergic agonists in hippocampus slices (Konopacki et al., 1987; Kowalczyk et al., 2013). Consistent with those reports, the action of acetylcholine in our study was not dependent on a rhythmic input since septal units decreased their discharge preference in the theta range during laser stimulation and the prolonged laser pulse did not entrain any particular rhythm.

It has been shown that the direct activation of septal cholinergic somata with laser trains under urethane anesthesia increases the power and synchrony of theta oscillations in the hippocampus (Vandecasteele et al., 2014; Dannenberg et al., 2015). The increased theta power arises partly from the suppression of peri-theta frequencies, thus resulting in an enhanced signal-to-noise ratio for the theta band. Conversely, we did not detect such changes in the peri-theta frequency bands during inhibition of somatostatin cells, but a very selective increase in the power of the theta band itself. In addition, previous results have shown that optogenetic stimulation of septal cholinergic cells dramatically suppressed ripple episodes in the hippocampus (Vandecasteele et al., 2014; Dannenberg et al., 2015), a result that we could not attain with the silencing of septal somatostatin cells, as ripple episodes were only partially, yet significantly decreased. Suppression of sharp-wave ripples might result from the direct activation of interneurons in the CA3 field (Dannenberg et al., 2015). Importantly, it has been proposed that cholinergic activation of theta oscillations and suppression of sharp-wave ripples rely on anatomically distinct pathways, with the indirect pathway being responsible for theta enhancement, and the direct pathway, which suppresses sharp-wave ripples (Dannenberg et al., 2015). In that case, our results would be better explained by the preferential activation of the indirect pathway, which boosts theta oscillations, and consistent with the differential connectivity between cholinergic and somatostatin neuronal populations.

During examination of the causal interactions between the medial septum and dorsal hippocampus, we found a significant unidirectional influence of the septum over the hippocampus. Urethane anesthesia induces prominent cortical slow waves, so it has been used as a model system to study sleep slow oscillations (Clement et al., 2008; Negrón-Oyarzo et al., 2018). During slow wave sleep, there is prominent unilateral causality in the septum to hippocampus direction in the theta frequency band (Kang et al., 2017a), which is similar to the interaction that we describe here under urethane anesthesia. Indeed, we found little descending influence from the hippocampus to the septum, but a prominent peak in theta activity in the opposite direction. Interestingly, despite the prominent theta peak in Granger causality, neither during slow wave sleep (Kang et al., 2017a) nor during our urethane-anesthesia experiments were theta oscillations evident in hippocampus field recordings. This observation has been interpreted as resulting from the little descending theta drive in the hippocampus to septum direction, which is likely necessary to reveal theta oscillations in the field potential (Kang et al., 2017a). Furthermore, intraseptal injections of somatostatin or the somatostatin receptor agonist octreotide in freely-moving rats reduced the power of hippocampal EEG in the theta band (Bassant et al., 2005), thus confirming that somatostatin cells participate in the control of hippocampal theta oscillations. Those results complement our findings, as they suggest that during somatostatin cell activation hippocampal theta waves are depressed, while our experiments show that during somatostatin cell inhibition hippocampal theta oscillations are enhanced. Interestingly, our results suggest that not only the power, but also the directionality of theta oscillations relies on the activity of somatostatin cells.

Arousal, locomotion, and theta oscillations are commonly concurrent, yet their mechanisms can be dissociated and independently controlled (Montgomery et al., 2009; Fuhrmann et al., 2015; Vinck et al., 2015). Importantly, the medial septum seems to be a neural hub for the regulation of all these processes. For example, it has recently been shown that medial septum glutamatergic neurons can disinhibit pyramidal cells in the hippocampus and control both theta oscillations and locomotion speed (Fuhrmann et al., 2015). Hence, goal-directed behavior, such as memory-guided spatial navigation, might be influenced by the action of somatostatin cells and their enhancement of theta oscillations in the dorsal hippocampus. On the other hand, septal somatostatin cells innervate the main local neuronal populations, yet they also express prominent descending projections reaching subcortical targets, like the lateral hypothalamus (Deng et al., 2019), that have been shown to control feeding preferences (Zhu et al., 2017) and fear conditioning behavior (Besnard et al., 2019). Indeed, septal somatostatin neurons can gate mobility to calibrate context-specific fear behavior, with descending contextual information provided by the CA3 area (Besnard et al., 2019). Importantly, septal somatostatin cells produce terminal fields that selectively target the lateral hypothalamus, where a distinct population of glutamatergic neurons powerfully controls the generation of repetitive self-grooming (Hong et al., 2014). Hence, a reasonable interpretation of our results is that silencing septal somatostatin disinhibits lateral hypothalamic glutamatergic cells, thus gating self-grooming behavior. This hypothesis will have to be tested in future experiments, yet it goes in line with recent data showing that septal somatostatin cells bidirectionally modulate conditioned fear responses by recruiting hypothalamic circuits during context discrimination. Indeed, optogenetic silencing of septal somatostatin cells increases contextual fear responses such as freezing behavior, also consistent with lateral septal units inversely tracking aversive conditioned stimuli (Besnard et al., 2019). Additionally, descending theta oscillations through the hippocampo-septal pathway regulate the speed of locomotion. Indeed, theta-patterned stimulation of lateral septum GABAergic projections to the lateral hypothalamus is sufficient to decrease ambulatory behavior (Bender et al., 2015). Overall, our results provide additional evidence supporting the role of septal somatostatin cells in the control of both motivated and innate behaviors.

Interestingly, human studies have confirmed a role for hippocampal theta oscillations in both spatial cognition and anxiety behavior (Harris and Gordon, 2015; Khemka et al., 2017). Moreover, while spatial navigation was associated with faster theta waves (4-8 Hz), threat-related anxiety was related to slower theta waves (2-6 Hz). Such distinction in frequency bands is reminiscent of the type 1 and 2 rodent theta bands (Cornwell et al., 2012). In recent years, a quantitative model based on the concept of velocity-controlled oscillators has been proposed for the septo-hippocampal circuit (Burgess, 2008), in which type 1 and type 2 theta oscillations determine distinct aspects of the overall theta frequency (Wells et al., 2013). Interestingly, despite their neurochemical differences, anxiolytic drugs reduce the immobility hippocampal theta frequency (Gray and McNaughton, 2008). Conversely, studies in rodents have shown that the power of theta oscillations, particularly in the ventral hippocampus, increases during anxiety-like behavior (Adhikari et al., 2010). These results suggest a tight relation between hippocampal theta oscillations and anxiety, and self-grooming is an extensively described mechanism for coping with anxiety in rodents (Kalueff et al., 2016). Our results show that regulating activity levels of septal somatostatin cells can affect both theta oscillations and self-grooming behavior. It has been previously suggested that monitoring self-grooming is a valuable measure of repetitive behaviors in preclinical models of neuropsychiatric disorders (Kalueff et al., 2016). Hence, we suggest that future investigations should consider septal somatostatin cells as a potential target for the control of altered innate behaviors in translational neuroscience (Negrón-Oyarzo et al., 2015).

## Acknowledgments

Grants Fondecyt 1190375, ANID/ACT210053

## Extended Data

**Figure 1-1.**
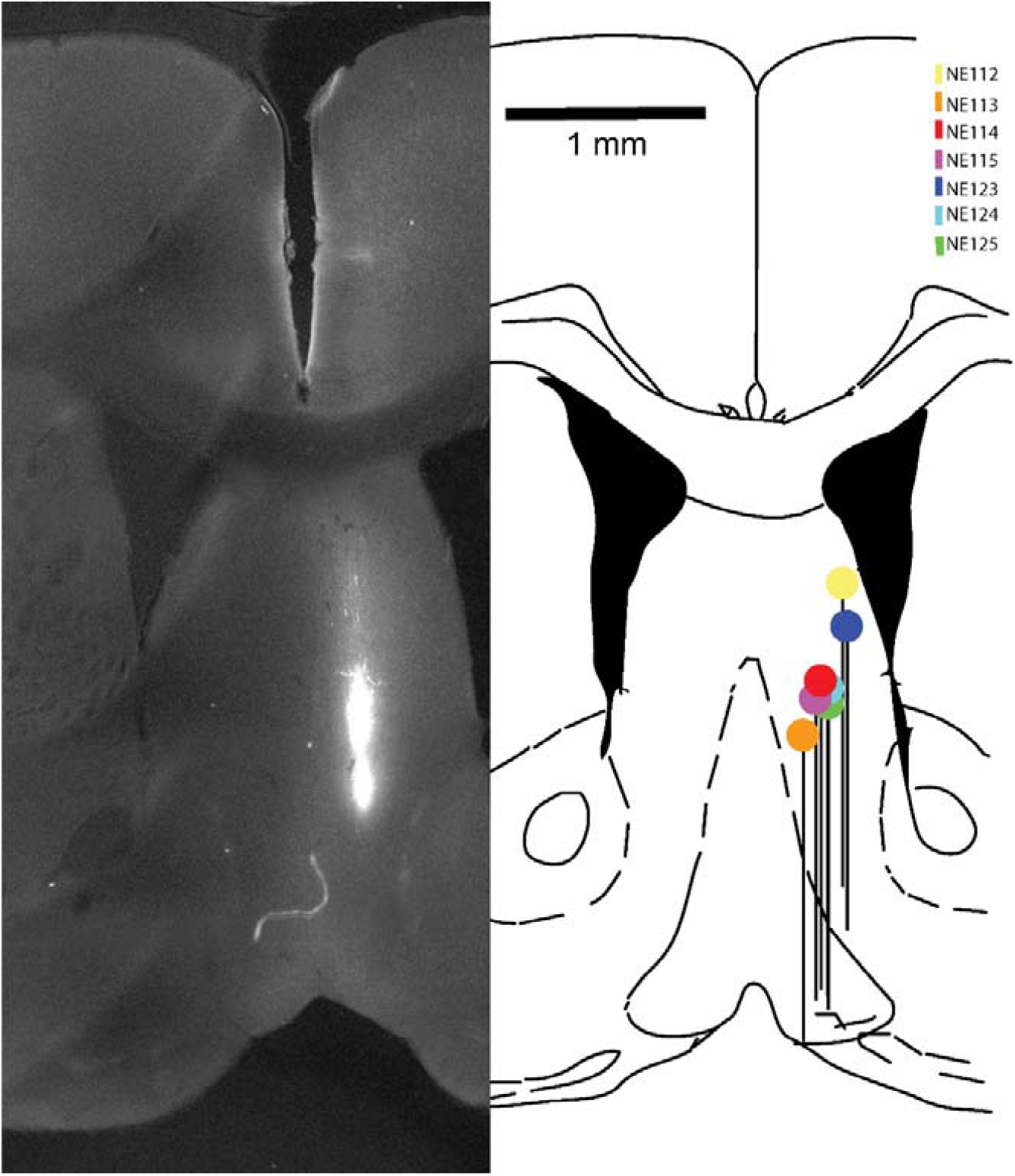
Extended Data figure supporting Figure 1. Anatomical location of recording sites. Left, example fluorescence microscopy image marking the placement of the tip of the fiber (NE 113). Right, representation of all experiments and placement of the tip of the fiber (colored circles) and the projection of the probe (black horizontal lines). Scale bar 1mm. Antero-posterior 0.98mm in reference to standard atlas coordinates.

**Figure 1-2.**
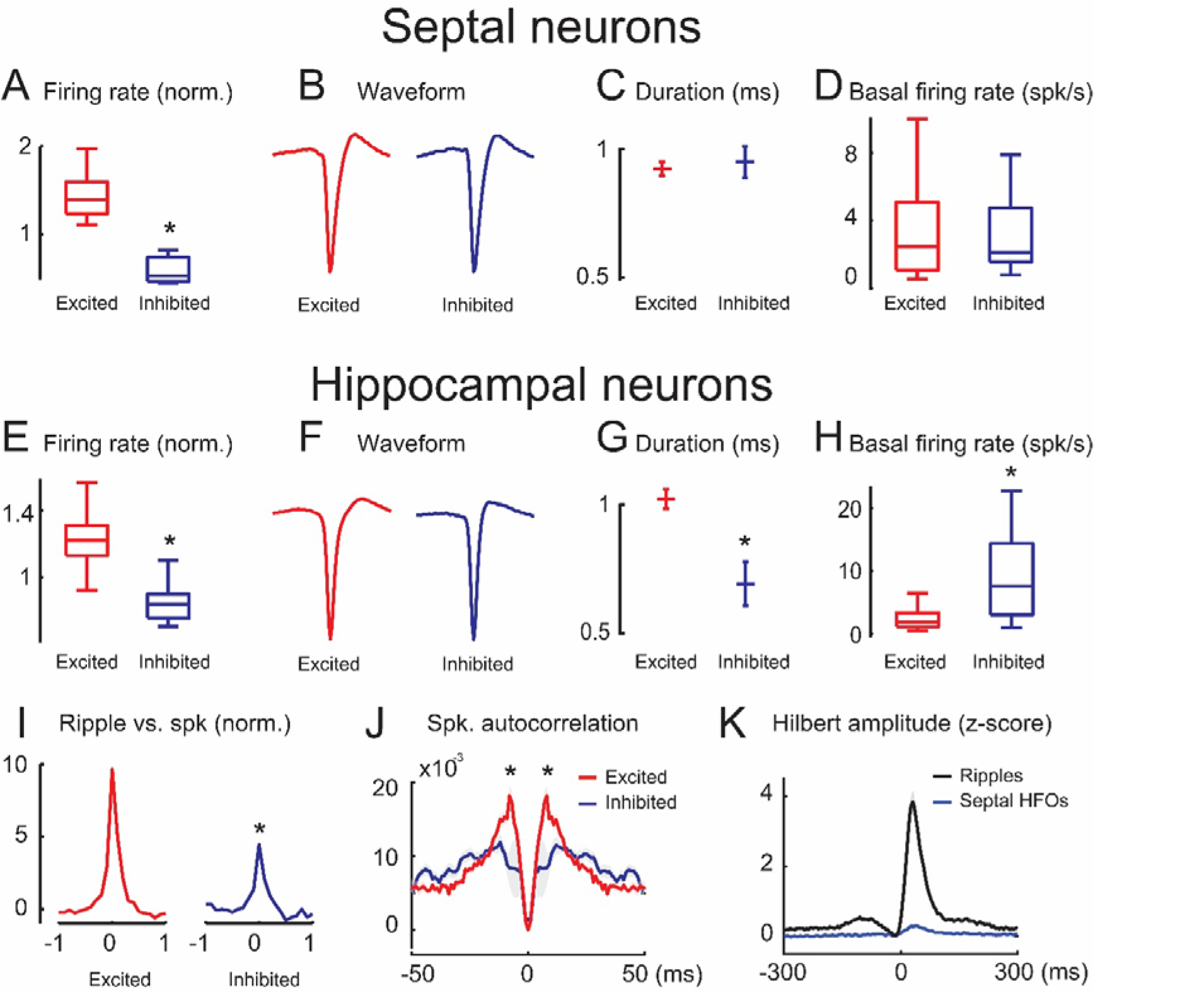
Extended Data figure supporting Figure 1. Neuronal activity in the medial septum and dorsal hippocampus during optical inhibition of septal somatostatin cells. Septal neurons: **(A)** normalized discharge probability during optical stimulation for excited (red line, n = 38) and inhibited (blue line, n = 13) neurons (Wilcoxon rank-sum test, *P = 9.94×10^−8^). **(B)** spike waveform average. **(C)** waveform peak-to-valley duration (Wilcoxon rank-sum test, P = 0.44). **(D)** basal firing rate (Wilcoxon rank-sum test, P = 0.47). Hippocampal neurons: **(E)** normalized discharge probability during optical stimulation for excited (red line, n = 65) and inhibited (blue line, n = 15) neurons (Wilcoxon rank-sum test, *P = 1.94×10^−9^). **(F)** spike waveform average. **(G)** waveform peak-to-valley duration (Wilcoxon rank-sum test, *P = 0.018). **(H)** basal firing rate (Wilcoxon rank-sum test, *P = 8.63×10–4). **(I)** crosscorrelogram of spiking activity related to hippocampal sharp wave ripples onset (unpaired t-test, *P = 0.0049). **(J)** spiking activity autocorrelation (Wilcoxon rank-sum test, *P = 0.032). **(K)** Ripple-triggered average between the ripple onset (reference) and the Hilbert amplitude for hippocampal LFP (100 – 250 Hz) and high frequency oscillations (HFOs) for Septal LFP (100 – 180 Hz), n = 18 recordings.

**Figure 2-1.**
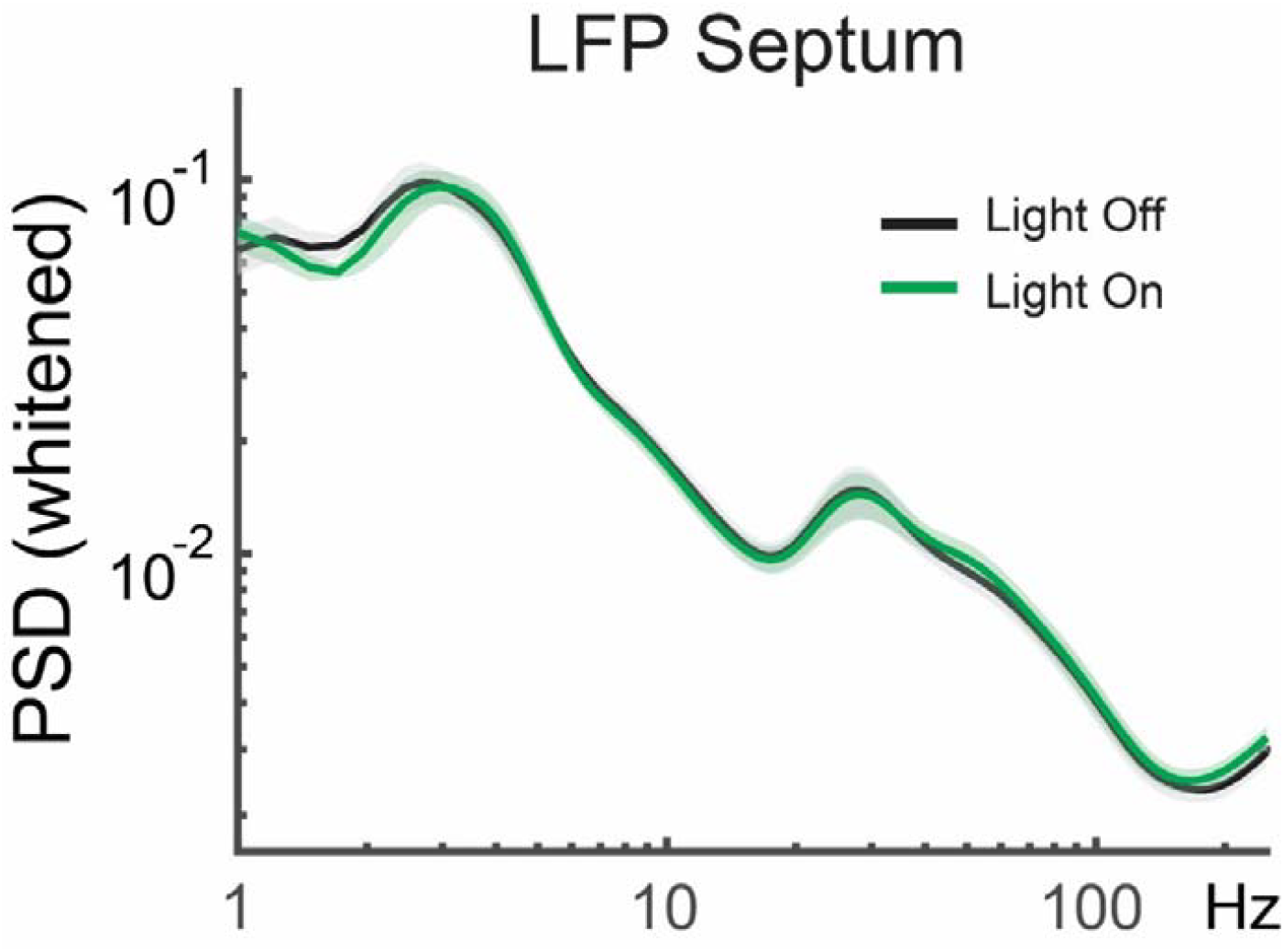
Extended Data figure supporting Figure 2. Average normalized power spectral density (PSD) of the septal LFP before (Control, black line, 5s) and during light stimulation (Light, green line, 5s)

**Figure 2-2.**
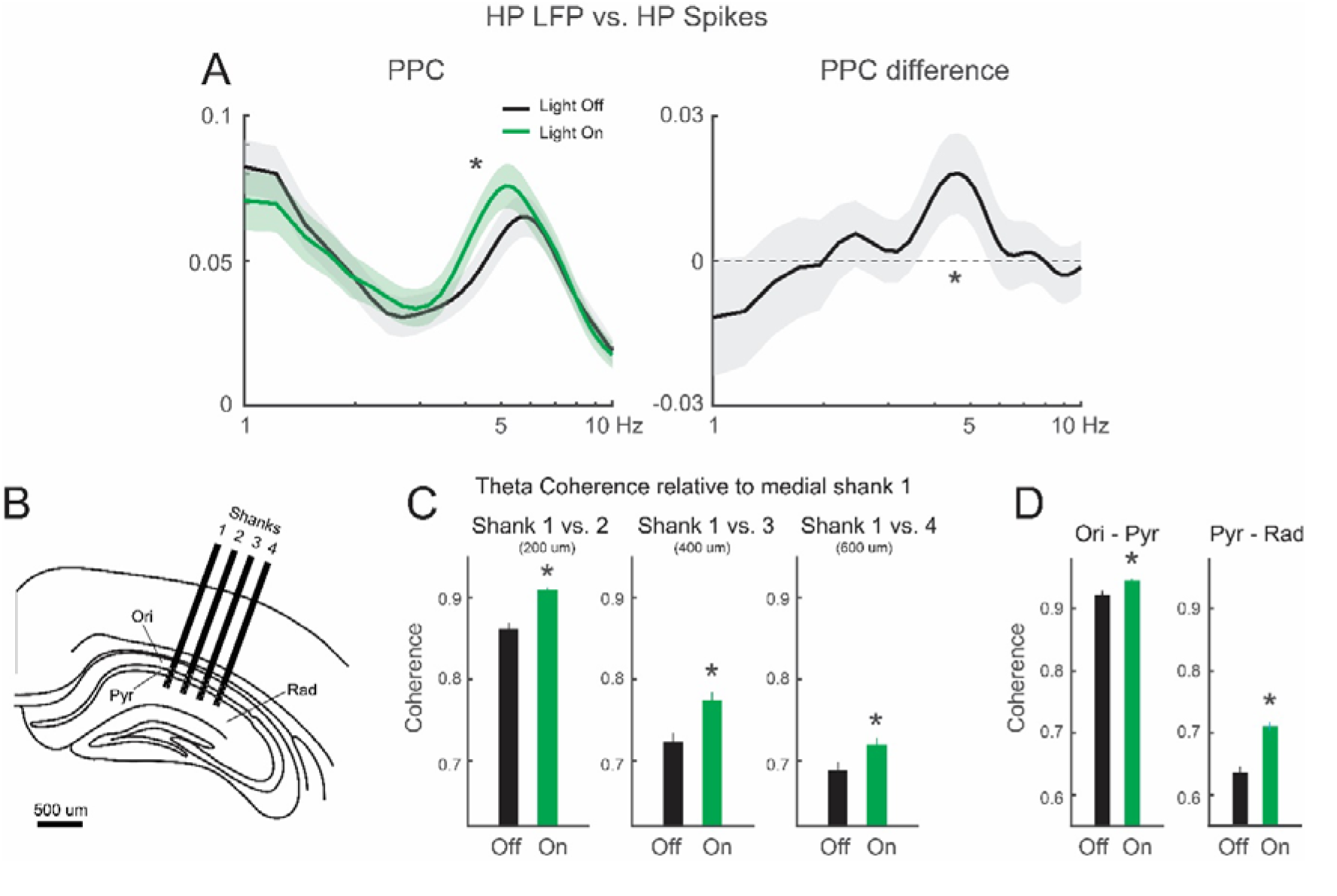
Extended Data figure supporting Figure 2. Intrahippocampal synchrony during optogenetic inhibition of septal somatostatin cells. **(A)** left: average pairwise phase consistency (PPC) in the dorsal hippocampus before (Control, black line, 5s) and during light stimulation (Light, green line, 5s) and, right: average PPC difference. Wilcoxon signed-rank test in band 4 – 6Hz. W = 21510, *p = 0.0189, n = 321 cells. **(B)** statistically significant increase of hippocampal theta coherence does not reveal interaction with the distance in the medio-lateral axis i.e., their magnitude is independent of the distance between shanks (two-way ANOVA, p ≈ 0). **(C)** the theta-promoting effect of septal stimulation was segregated between hippocampal layers, being more prominent between stratum radiatum and pyramidal layer. Oriens – pyramidal coherence: Wilcoxon signed-rank test, W = 2904, *p = 2.24×10^−5^, n = 140 trials. Pyramidal – stratum radiatum: Wilcoxon signed-rank test, W = 1213, *p = 3.65×10^−29^, n = 210 trials.

**Figure 3-1.**
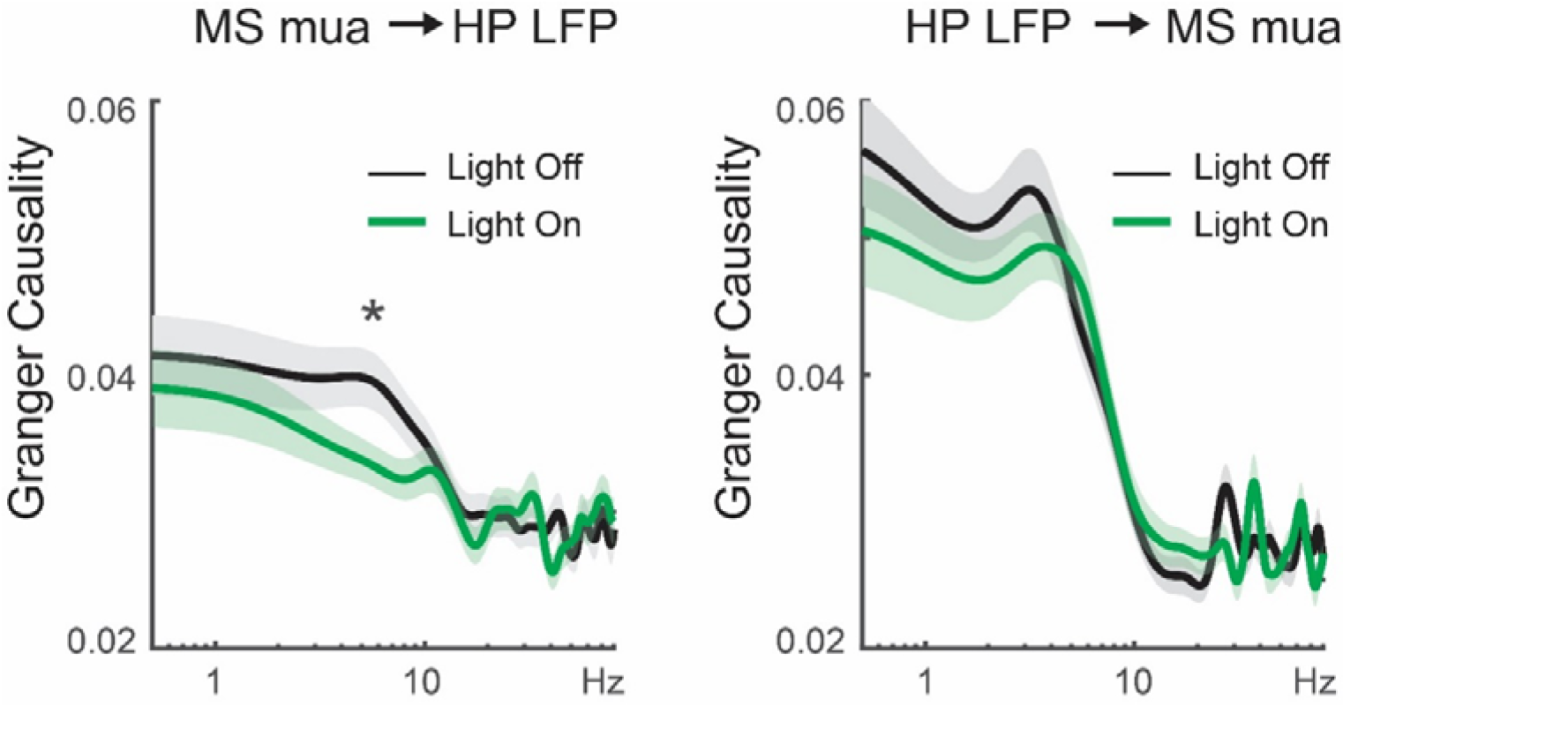
Extended Data figure supporting Figure 3. Effect of dorsal septum optogenetic stimulation in Granger causality between septal multiunit activity and hippocampal LFP. **(A)** MS->HP connectivity in theta band was weakened by inhibition of septal somatostatin neurons. Asterisk in band 3 – 7 Hz: paired t-test, p = 0.0143, n = 405 trials. **(B)** HP->MS connectivity is apparently unaffected by optogenetic stimulation (but see main text and Fig. 3 for details).

**Figure 4-1.**
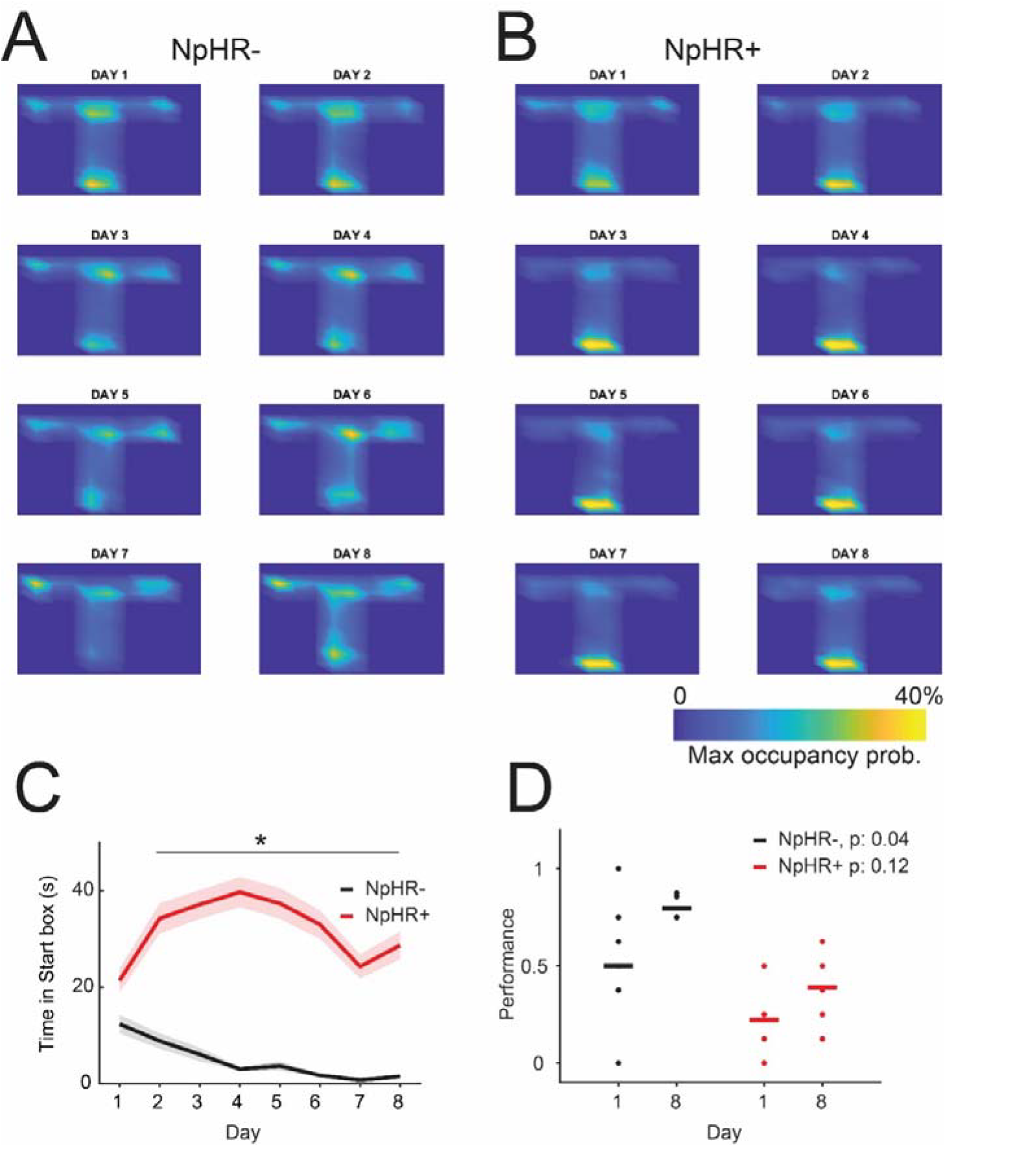
Extended Data figure supporting Figure 4. Average normalized occupancy at each location within the T-maze through the days for **(A)** NpHR- and **(B)** NpHR+ animals. **(C)** time in the start area (defined as the first 10 cms of the central arm where the animal is placed at the beginning of every trial). *p < 0.009, Wilcoxon rank sum test corrected by false-discovery rate. Values in mean ± SEM. **(D)** comparison of performance measurement between day 1 and day 8 for NpHR-(black, n = 8) and NpHR+ (red, n = 9) animals. Line denotes mean and dots the individual values. Respective differences were determined by a paired-sample t-test.

**Figure 4-2.**
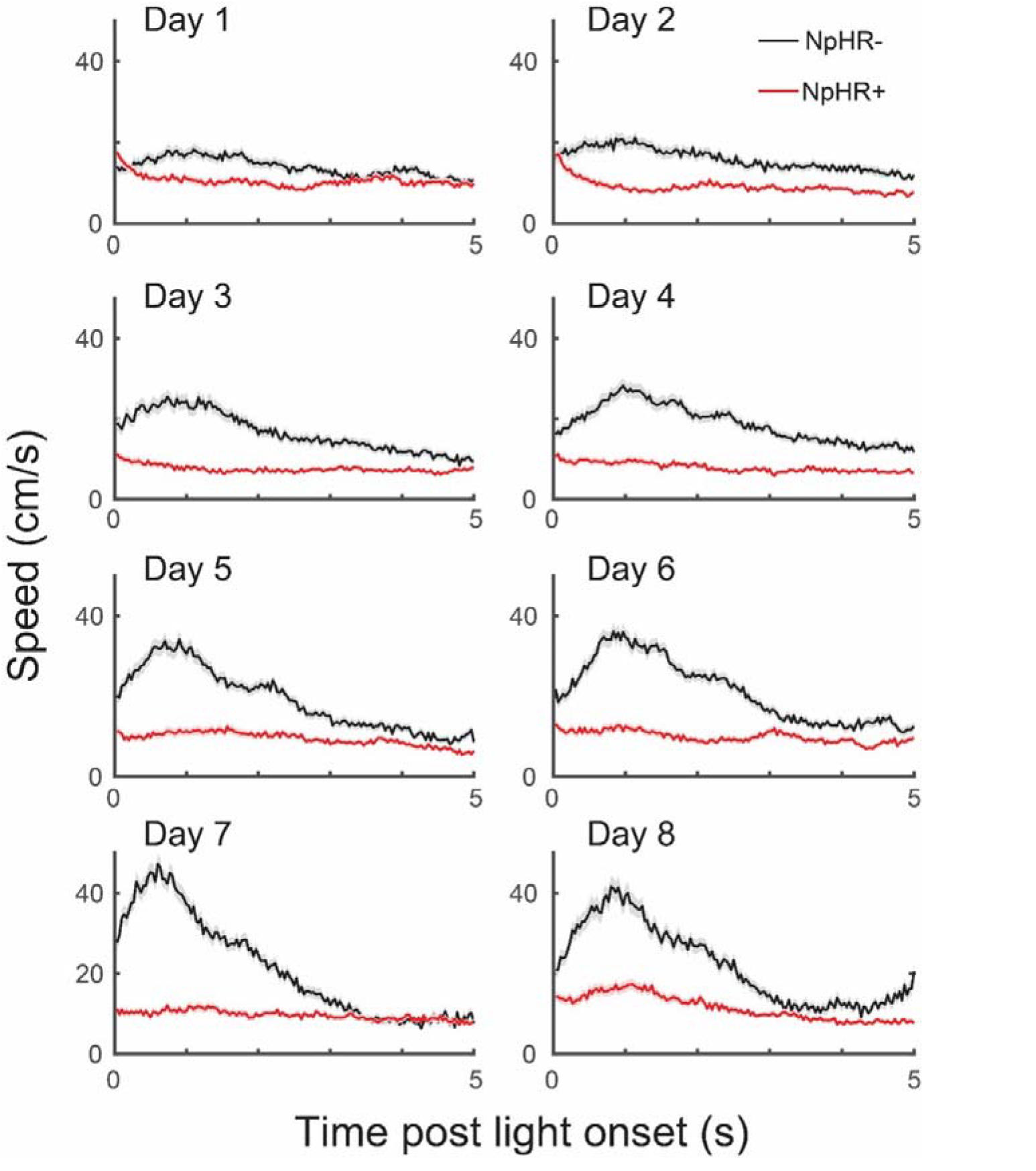
Extended Data figure supporting Figure 4. Average instantaneous speed for all trials in the T-maze through the days for NpHR-(black line) and NpHR+ (red line). Values in mean ± SEM. Bin = 33 ms.

**Figure 4-3.**
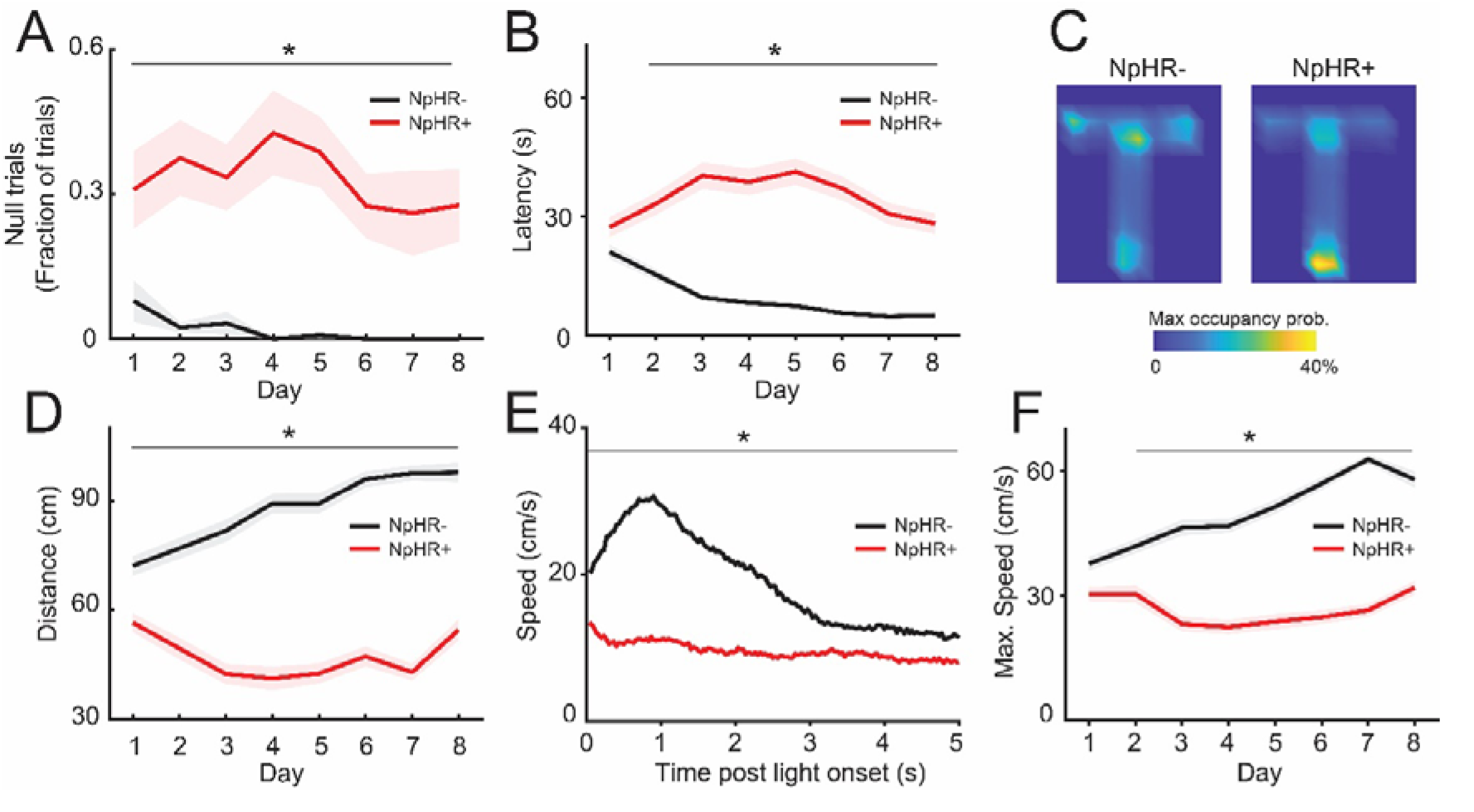
Extended Data figure supporting Figure 4. Behavioral parameters for non-null trials. **(A)** fraction of null trials for NpHR- and NpHR+ animals. Task latency **(B)**, average normalized occupancy **(C)**, total traveled distance **(D)**, average instantaneous speed during light stimulation **(E)** and, average maximum instantaneous speed **(F)** across days, represented for trials where animals effectively reached or consumed the reward within 90 s, i.e., non-null trials. Values in mean ± SEM; NpHR-, black line. NpHR+, red line. *p: significant pairwise comparison across days corrected by false-discovery rate. For (A) *p < 0.022, (B) *p < 0.023, (D) *p < 3×10^−18^, (E) *p < 3×10^−5^ and (F) *p < 0.001.

**Figure 4-4.**
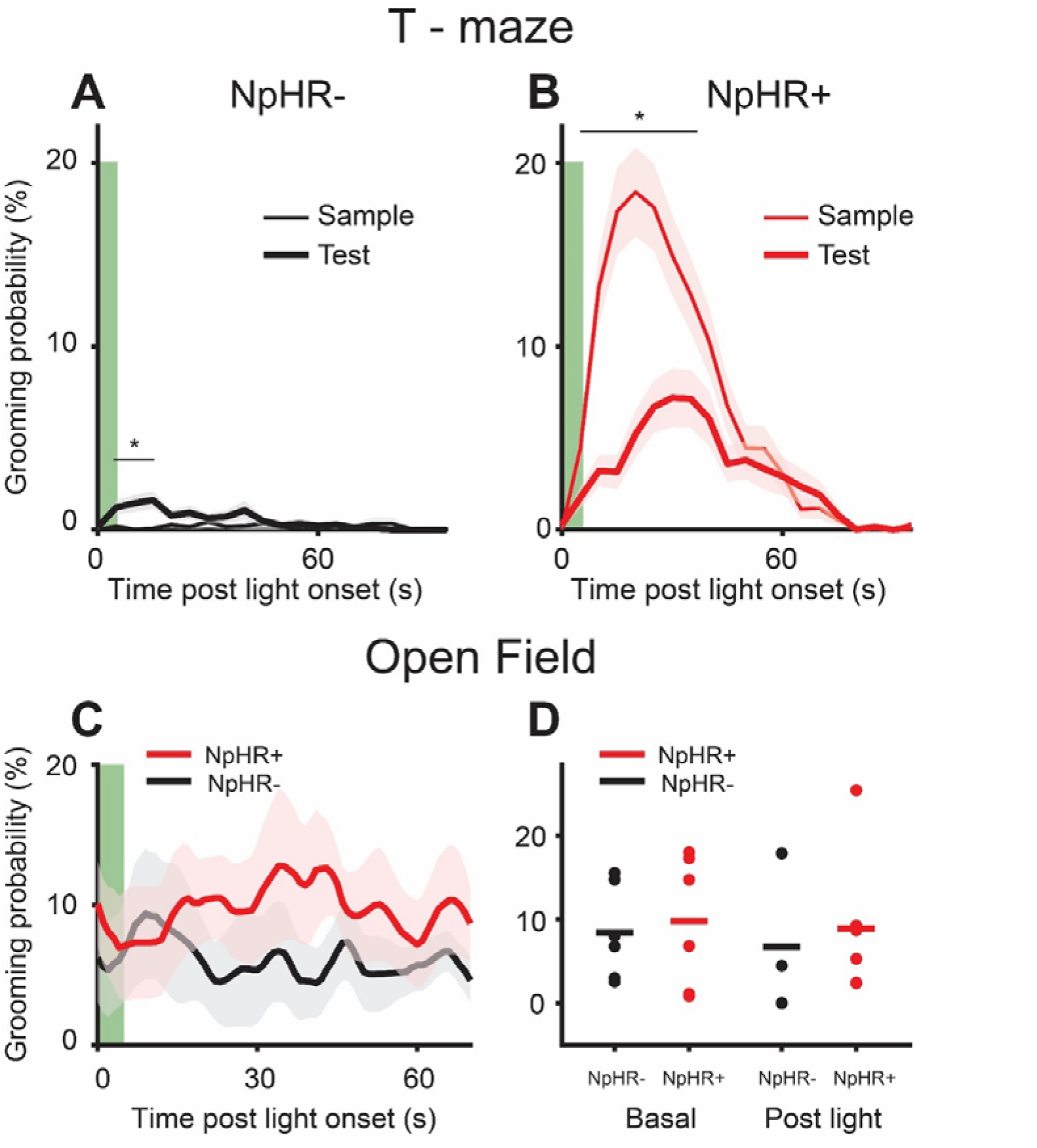
Extended Data figure supporting Figure 4. Average probability of grooming during a goal-directed behavior vs. a spontaneous open field exploration. **(A)** grooming during sample (thin line) and test (thick line) trials in the T-maze for NpHR-animals. *p < 0.0028, bin = 5 s. **(B)** grooming during sample (thin line) and test (thick line) trials in the T-maze for NpHR+ animals. *p < 0.0027, Bin = 5 s. **(C)** average probability of grooming in an open field after light stimulation for NpHR-(70 trials, n = 6 mice) and NpHR+ (69 trials, n = 6 mice) animals. Bin = 1 s. **(D)** two-way repeated-measures ANOVA showed no significant difference in averaged grooming probability both for basal condition and after light stimulation between NpHR- and NpHR+ animals. Green bar indicates light stimulation. Values in mean ± SEM. *P, significant pairwise comparison corrected by FDR.

**Table 2-1.** Extended Data table supporting Figure 2. Changes induced by light in hippocampal LFP rhythms.

| Frequency band | Light Off | Light On | p |
| --- | --- | --- | --- |
| Slow band (0.5 - 3 Hz) | 73.2 ± 3.3 | 71.9 ± 3.4 | 0.531 |
| Theta (4 - 7 Hz) | 68.3 ± 3.3 | 78.4 ± 4.4 | 7.99x10<sup>-7</sup> (*) |
| Slow-Gamma (20 - 40 Hz) | 22.5 ± 1.9 | 23.1 ± 1.8 | 0.266 |
| High-gamma (60 - 140 Hz) | 6.6 ± 0.5 | 6.8 ± 0.5 | 0.158 |

Values in mean ± SEM 10<sup>-3</sup> x (uV<sup>2</sup>/Hz). p = paired t-test, asterisk denotes statistical significance.

**Table 2-2.**
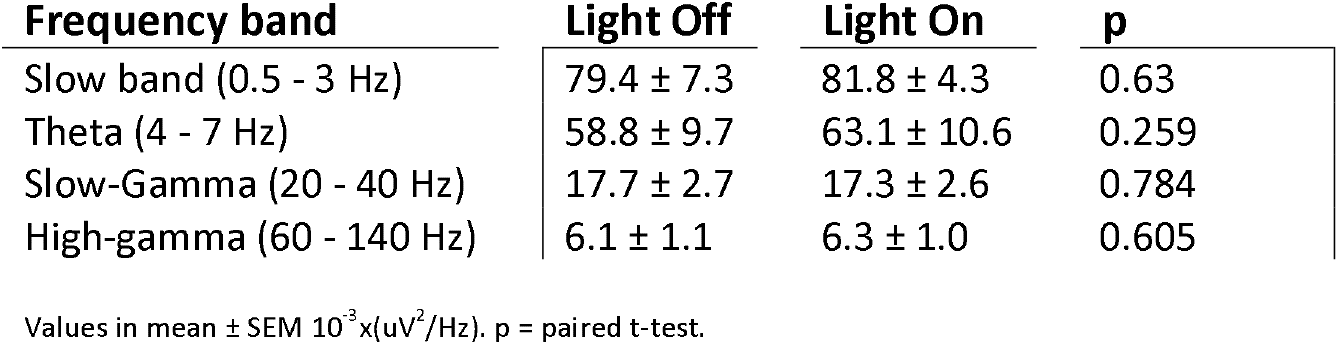
Extended Data table supporting Figure 2. Changes induced by light in hippocampal LFP rhythms under ACh receptors blockade.

**Table 3-1.** Extended Data table supporting Figure 3. Changes induced by light on Granger causality from the septal LFP to hippocampal LFP (MS LFP -> HP LFP).

| Frequency band | Light Off | Light On | p |
| --- | --- | --- | --- |
| Slow band (0.5 - 3 Hz) | 0.16 ± 0.009 | 0.15 ± 0.008 | 0.97 |
| Theta (3 - 6 Hz) | 0.18 ± 0.008 | 0.15 ± 0.007 | 2.4x10<sup>-6</sup> (*) |
| Slow-Gamma (20 - 40 Hz) | 0.08 ± 0.003 | 0.075 ± 0.003 | 0.27 |
| High-gamma (60 - 140 Hz) | 0.06 ± 0.001 | 0.06 ± 0.001 | 0.74 |

Values in mean ± SEM. p = Wilcoxon signed-rank test.

**Table 4-1.**
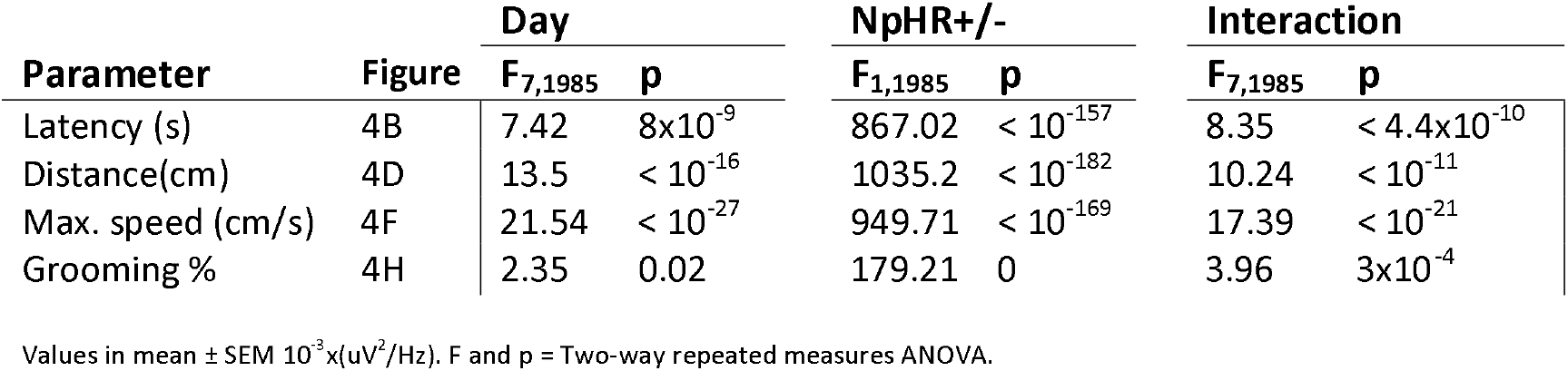
Extended Data table supporting Figure 4. Descriptive statistics for behavioral data in a spatial memory task.

| Parameter | Figure | Day |  | NpHR+/- |  | Interaction |  |
| --- | --- | --- | --- | --- | --- | --- | --- |
|  |  | F<sub>7,1985</sub> | p | F<sub>1,1985</sub> | p | F<sub>7,1985</sub> | p |
| Latency (s) | 4B | 7.42 | 8x10<sup>-9</sup> | 867.02 | < 10<sup>-157</sup> | 8.35 | < 4.4x10<sup>-10</sup> |
| Distance(cm) | 4D | 13.5 | < 10<sup>-16</sup> | 1035.2 | < 10<sup>-182</sup> | 10.24 | < 10<sup>-11</sup> |
| Max. speed (cm/s) | 4F | 21.54 | < 10<sup>-27</sup> | 949.71 | < 10<sup>-169</sup> | 17.39 | < 10<sup>-21</sup> |
| Grooming % | 4H | 2.35 | 0.02 | 179.21 | 0 | 3.96 | 3x10<sup>-4</sup> |

Values in mean ± SEM 10<sup>-3</sup> x (uV<sup>2</sup>/Hz). F and p = Two-way repeated measures ANOVA.

